# Multiparental RNA-seq driven eQTL screening identifies loci underlying host plant fitness in a generalist herbivore

**DOI:** 10.64898/2026.08.30.748081

**Authors:** Femke De Graeve, Berdien De Beer, Sander De Rouck, Shauny Naessens, Mathijs Herpoel, Jan Fostier, Richard M. Clark, Thomas Van Leeuwen, Tim De Meyer

## Abstract

The two-spotted spider mite (*Tetranychus urticae*) is an extremely polyphagous pest, yet the genetic basis of this adaptive potential remains to be fully elucidated. Since expression quantitative trait loci (eQTLs) provide the genetic basis of numerous phenotypes, we aimed to identify *trans*-eQTL hotspots underlying *T. urticae* fitness upon transfer from a common (bean) to a challenging (tomato) host plant. Nonetheless, the identification of *trans*-eQTLs is complex and often constrained by methodological challenges and high costs. Therefore, we employed a multiparental mapping strategy driven by RNA-seq, enabling us to leverage extensive genetic variation in a cost-efficient manner. A randomly mating population was generated on bean from a small number of genetically diverse, often heterozygous parents, and subsequently transferred to tomato prior to RNA-seq. Upon whole genome sequencing of the parents, RNA-seq of the mapping population individuals was sufficient to reconstruct their genomes as a combination of parental haploblocks. Subsequent eQTL mapping identified 23 distinct *trans*-eQTL hotspot regions associated with the expression of numerous target genes. The most prominent hotspot on chromosome 3 was associated with approximately 900 genes and showed enrichment for functions related to detoxification and digestion. Furthermore, 4 of these *trans*-eQTL hotspot genotypes explained significant variation in mite fitness on the challenging host plant. This contrasted a traditional QTL mapping approach, where these genotypes could not be detected due to multiple testing correction. Our study hence offers a powerful strategy for *trans*-eQTL hotspot screening and the discovery of trait-associated loci in complex, multiparental genetic backgrounds.

## 1. Background

Quantitative Trait Locus (QTL) studies are fundamental for identifying the genetic basis of complex traits by locating genomic regions associated with phenotypic variation^1^. An important class of QTLs are Expression Quantitative Trait Loci (eQTLs), genetic variants that affect gene expression levels^2^. Acting as a bridge between genotypes and phenotypes, eQTL studies provide crucial insight into how genetic variation impacts phenotypic traits through altered gene expression, and are essential for inferring gene regulatory networks^3–6^. A key distinction is made between *cis-* and *trans*-eQTLs. *Cis-* eQTLs are located close to the target gene and influence expression in an allele-specific manner, while *trans*-eQTLs are typically located further away and affect both alleles of the target gene^3,7,8^. In particular, *trans*-regulatory hotspots, where a single locus affects multiple genes, are of interest as they coordinate gene expression patterns and can inform systems-level understandings of phenotypic variation^9–11^. The effect of a *trans*-hotspot variant can be direct, e.g. when transcription factor variation affects binding affinity to its target genes, and/or indirect, when a variant leads to an altered physiology with associated expression changes. In both cases, however, the variant’s impact on many genes makes it a prime candidate to explain phenotypic variation.

In practice, eQTL mapping studies face important technical and budgetary challenges, limiting their success. Typically, these studies require both genotype and expression data for all individuals in a large mapping population, leading to expensive designs^5^. A common alternative, when practically and ethically feasible, involves the generation of inbred lines for crossing and subsequent sequencing of the progeny, which can maximize power while minimizing the number of individuals that need to be sequenced at the DNA and RNA level. The use of inbred lines also simplifies genotyping, as their homozygosity reduces ambiguity in allelic inheritance. Yet, generating inbred lines involves significant effort, potential challenges in maintaining them due to reduced fitness^12,13^, and inherently limits the genetic variability under study. Further, from a statistical perspective, eQTL mapping is not trivial. Detecting *trans*-eQTLs is particularly challenging because of their generally smaller effect sizes, a larger tissue-specificity, and the number of statistical tests^3,10,14^. Indeed, testing for associations between the expression of all genes and all single nucleotide polymorphisms (SNPs) typically results in millions, if not billions, of statistical tests, significantly reducing statistical power owing to the extreme multiple testing burden^14^.

In this study, we address common challenges in eQTL mapping with an agriculturally relevant proof-of-concept study examining the transcriptional response of *Tetranychus urticae*, the two-spotted spider mite, to a novel host plant. This mite is an excellent use case for studying the genetic basis of transcriptional regulation due to its exceptional ability to develop pesticide resistance and quickly adapt to new host plants^15,16^. Moreover, this haplodiploid organism has a well-annotated genome assembly^17,18^.

Previous research by Kurlovs et al.^19^ highlighted extensive *trans*-regulation in *T. urticae*, particularly of detoxification genes, such as cytochrome P450 monooxygenases (P450s), carboxyl-choline esterases (CCEs), ATP-binding cassette (ABC) transporters, major facilitator superfamily (MFS) transporters, intradiol-ring cleavage dioxygenases (DOGs), UDP-glucuronosyltransferases (UGTs) and lipocalins, which play critical roles in pesticide resistance and host plant adaptation. Similarly, Ji et al.^11^ identified a *trans*-eQTL hotspot in a pesticide resistant spider mite strain regulating a large set of detoxification genes. Collectively, these studies suggested a central role of eQTLs in transcriptional regulation of detoxification. Nevertheless, it remains unclear which particular *trans* effects underlie specific adaptive phenotypes in this species. Moreover, the need for complex and experimentally challenging crossing schemes and sample collection strategies limits the approaches used by Kurlovs et al.^19^ and Ji et al.^11^ for studies within and beyond spider mite.

Therefore, in this manuscript, we introduce a cost-efficient method for eQTL mapping and illustrate it for *T. urticae* using fecundity on a new host plant as the phenotype of interest. The rationale is that whenever a small number of genetically diverse diploid individuals can be used as founding parents to create an artificial randomly mating population, only whole-genome sequencing (WGS) of the founding parents and RNA-sequencing (RNA-seq) data of the mapping population individuals are required for eQTL mapping. Indeed, RNA-seq of the mapping population individuals is sufficient to reconstruct their genomes as recombinations of the WGS inferred founding parental genomes, thereby yielding haploblock-based resolution. Genetically determined gene expression is subsequently inferred per gene by means of regularized regression, leading to the identification of *cis*- and *trans*-eQTLs. This strategy not only mitigates the presence of RNA specific or other artefacts at the SNP/gene level, but also enhances the power for *trans*-eQTL detection by markedly reducing the multiple testing burden. Moreover, it is flexible in the sense that the founding parents of a “*quasi*-panmictic” population may be heterozygous as well as (partially) homozygous.

Based on the eQTL mapping, we identified 23 distinct *trans*-eQTL hotspots. A major hotspot on chromosome 3 was associated with approximately 900 genes and showed enrichment for functions related to detoxification of plant-derived metabolites and digestion. Subsequently, eQTLs relevant for the phenotype of interest – fitness on tomato – were identified by regressing genetically determined gene expression on mite fecundity on the new host. Finally, we demonstrate that fitness on an alternative host in spider mites is (at least partially) mediated by genomic intervals harboring *trans*-eQTL hotspots, thereby linking gene expression patterns to organismal fitness. In summary, our study introduces a generic eQTL screening strategy generally applicable to any setting where a *quasi*-panmictic population can be easily generated from a small number of genetically diverse parental founders.

## 2. Results

In the results section, we first outline the general methodology (2.1), with the technical details of the procedures described in the corresponding Methods section. Subsequently, we elaborate on the design and basic characteristics of the study (2.2), followed by genotyping of the generated *quasi*-panmictic mapping population (2.3). Thereafter we screen for eQTLs (2.4) and identify those *trans-*eQTL hotspots associated with fitness on the new host, tomato (2.5).

### 2.1 Overview of the methodology

The rationale of the methodology is that in a multiparental, *quasi*-panmictic mapping population created from genetically diverse, founding parents, one can reconstruct the mapping population’s individual genomes as mosaics of the parental genomes. A concept that has been used in QTL mapping studies using multiparental populations – including heterogeneous stock^20^ and diversity outbred^21,22^ mice populations, and multiparent advanced generation intercross (MAGIC) populations in plants^23,24^. Unlike the more traditional designs, our approach does not rely on fully inbred or phased parental haplotypes, uses RNA-seq to genotype the mapping population, and applies a haploblock-based framework to mitigate alignment artefacts, including those caused by extensive copy number variation (CNV) and cross-mapping of short reads. Henceforth, mapping population individuals and founding parents will be referred to as “individuals” and “parents”, respectively.

Our decomposition methodology starts with obtaining WGS-based parental genotypes (Figure 1A) and RNA-seq-based genotype probabilities for each SNP and individual (Figure 1B). The use of genetically diverse parents entails the presence of a large number of SNPs that discriminate between them. Hence, relying on loci homozygous within each parent (but variable between at least two parents), a Hidden Markov Model (HMM) is used to identify the parental origin across the individuals’ genomes (Figure 1C), simultaneously identifying the locations of inter-parental recombination events. Note that the HMM step alone would suffice to delineate haploblocks if the parents were entirely homozygous, or if individual parental genomes with fully phased haplotypes were available (e.g. from long-read sequencing), rather than population-level parental data (as is intrinsic to our study on minute spider mites). To account for this intra-parental genetic variation arising from the use of heterozygous (i.e. non-inbred) parents, de Bruijn graphs (DBGs) are constructed from the parental DNA data for allele assembly per gene and parent. Consequently, for each gene per individual, the RNA-based genotype can be matched to compatible parental allele combinations, i.e. all diploid combinations of DBG inferred alleles for the HMM inferred parents for that individual’s gene. The best matching parental allele combination is hence considered the initial genotype for that gene and individual.

**Figure 1.**
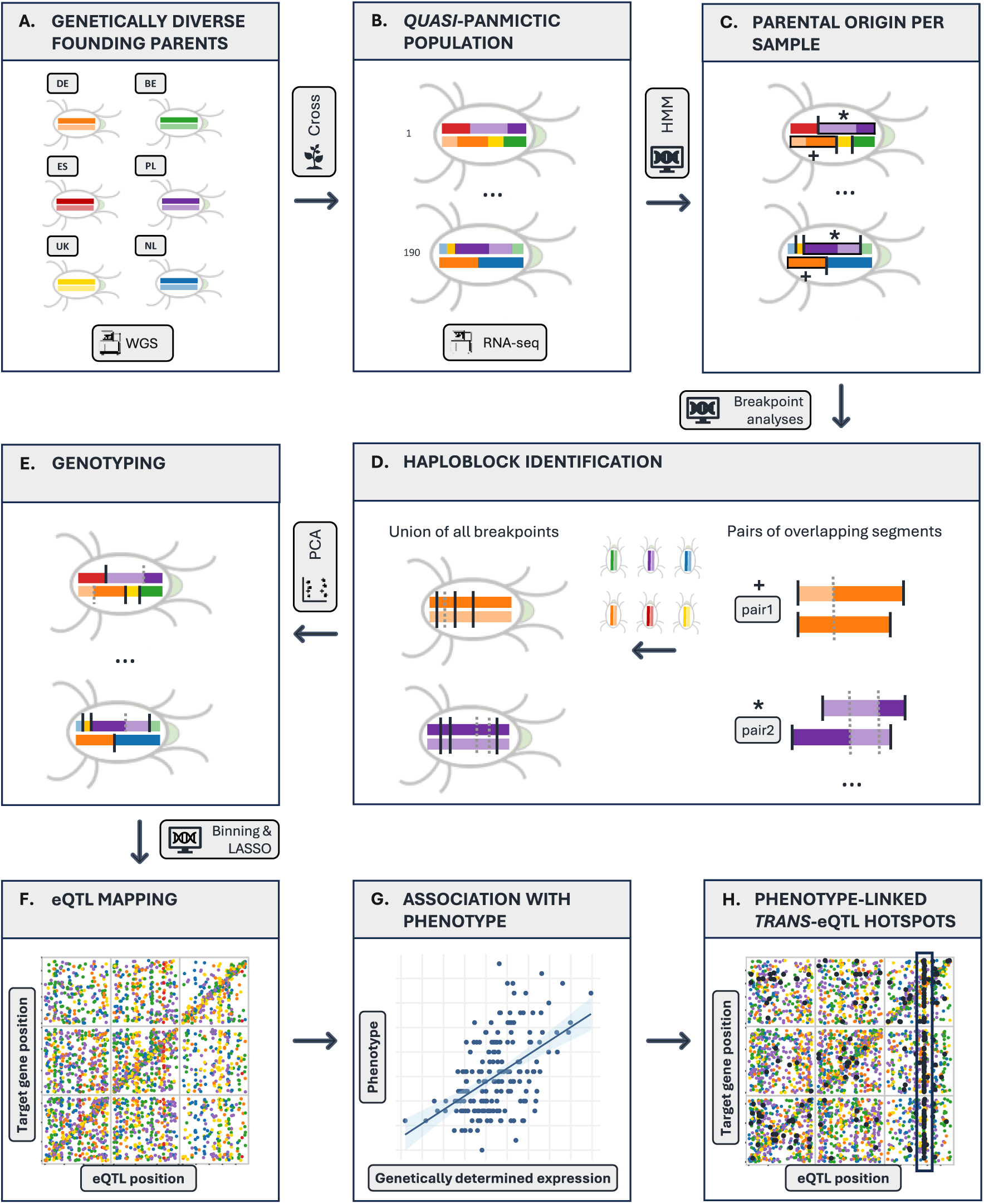
Schematic overview of the methodology based on a *T. urticae* case study. A) Six iso-female founders with WGS data were crossed to create B) a *quasi*-panmictic mapping population from which 190 individuals were randomly picked for RNA-sequencing. C) Homozygous SNPs per parental line were inferred from the WGS data and used as input of a Hidden Markov Model (HMM) to characterize the parental origin of the mapping population samples (with RNA-seq inferred genotype probabilities) along the genome and to identify inter-parental recombination events (black lines). D) Additional breakpoint analysis was used to delineate intra-parental recombination events (grey dotted lines) for (partially) heterozygous parents. We identified those breakpoints by comparing pairs of overlapping segments from the same parent in the mapping population individuals, yielding the final set of haploblocks. E) Per sample, PCA was used for genotyping at the haploblock level. F) Genotypes were compiled and made comparable (binning) for eQTL mapping by regularized regression. G) eQTLs associated with phenotype were located through the use of genetic instruments, i.e. genetically determined gene expression and, H) *trans* regulatory hotspots associated with the phenotype of interest identified.

These initial gene level genotypes are subsequently merged to form haploblocks with linkage information. This step mitigates RNA-seq artefacts at the gene level, e.g. when an allele was present at the DNA level but not (sufficiently) expressed, while also drastically reducing the multiple testing burden for later eQTL screening. The edges of haploblocks are determined by inter-parental recombination (Figure 1C), but also by the intra-parental recombination events (Figure 1D) that took place prior to or during the creation of the mapping population. In the *quasi*-panmictic mapping population, such intra-parental recombination may occur when – due to random chance – an individual features both alleles from the same parent in a certain genomic region.

Hence, to delineate haploblocks, intra-parental recombination events need to be identified per parental genome. Here, the non-informative homozygous regions in parents (if present) are ignored (see Methods section 5.3.4 for details). Using the HMM results, we identify genomic regions in individuals that include one allele of the parent under consideration and a second allele from any other parent. Crucially, in such a region, all variants specific for the parent under consideration are automatically linked, and hence represent a single DNA segment originating from that parent. By collecting and comparing all these DNA segments per parent, intra-parental recombination events can be identified. Specifically, per parent, we consider each pair of thus obtained DNA segments overlapping in the genome (e.g. pair 1 and 2 in Figure 1D) and perform breakpoint analysis using the initial gene-level genotypes. Resulting breakpoints represent those positions where overlapping DNA segments switch from predominantly sharing the same parental variant to predominantly featuring the other parental variant (or v.v.). The method can hence be made robust to initial gene-level genotyping errors (see Methods section 5.3.4 and 5.3.5 for details), yielding breakpoints that define haploblock boundaries.

Upon inferring the haplotypes per parental haploblock, each mapping population sample can subsequently be genotyped by matching the initial gene-level genotypes to the most likely haplotype per relevant haploblock, relying on principal component analysis (PCA) to remove inconsistent (hence likely aberrant) gene-level genotypes (Figure 1E). Ultimately, genotype bins are identified based on the union of all haploblock boundaries (including the HMM inferred inter-parental recombinations), so that these bins correspond to genomic regions featuring homogenous genotypes for all samples. Subsequently, variables are created for all parental haplotypes per bin. Each variable represents the dosage of a specific parental haplotype (encoded as 0, 1, or 2 to indicate the number of copies carried by an individual) and serves as a predictor in the LASSO regression for eQTL screening (see Supplementary Figure 1 for details on these genotype variables).

Per gene, regularized regression (Least Absolute Shrinkage and Selection Operator, LASSO) is then performed to predict gene expression based on the different bin genotypes (Figure 1F), enabling the discrimination of local effects (*cis*-eQTLs) from the impact of distant genotypes (*trans*-eQTLs). Finally, a genetic instrument strategy is used to identify eQTLs relevant to the phenotype of interest. Specifically, it is assessed whether genetically determined gene expression (LASSO prediction) is significantly associated with phenotype (Figure 1G). Finally, *trans*-eQTL hotspots significantly enriched for targets associated with the phenotype of interest are identified by overrepresentation analysis (Figure 1H). A technical description of the implementation is provided in the Methods section.

### 2.2 Design and baseline characteristics

To provide a proof of concept for this new strategy of eQTL mapping, a multiparental mapping population was created starting from 6 genetically diverse single female individuals (designated DE, BE, ES, PL, UK and NL based on their origin), as described earlier^25^. The phenotype under study was fitness on an alternative host, which we evaluated by migrating mites from an easy host on which nearly all *T. urticae* strains thrive (bean, *Phaseolus vulgaris*) to a challenging host plant (tomato, *Solanum lycopersum*). Consistent with previous research, fecundity (number of eggs laid) was used as a measure of fitness^25^. Gene expression was profiled on tomato, where *T. urticae* is known to exhibit a strong transcriptional response to host switch^25–29^.

Each single female had been expanded into an iso-female *T. urticae* line to facilitate genotyping of the parental populations through WGS, which is required given the minute size of individual mites. Since *T. urticae* is haplodiploid, unfertilized eggs of a single, virgin diploid female lead to haploid males, which can mate with their mother to create an iso-female line. *De novo* mutations aside, each line may hence feature at most two different haplotypes per locus, with drift leading to increasing homozygosity over time. Note that the use of iso-female lines has no impact on the generic methodology, as any recombination occurring within a line prior to the creation of the *quasi*-panmictic population will be detected as intra-parental recombination and included in the haploblock boundaries. In preliminary experiments, random mixing of individuals of the 6 iso-female lines did not lead to the expected random mating behaviour, as the resulting population (after 3 generations) still contained individuals that were genetically fully derived from a single parent (3 out of 5 individuals evaluated). Therefore, the mapping population used in this study was created by a 2-step inter-strain crossing scheme depicted in Villacis-Perez et al.^25^ and further expanded by random mating for ∼11 weeks (about 6 generations after the two initial crosses; throughout, bean was used as the host plant).

Subsequently, fecundity of individual females of the mapping population was assessed at three timepoints: (1) after 24 hours on their initial host, bean, (2) after 24 hours on the new host, tomato, and (3) after 48 hours on the new host (leaf enclosures with single females were used). Immediately afterwards, the female mites were frozen for individual RNA-isolation. Eggs were counted using machine learning driven image analysis of leaf disk photographs (performed with YOLOv5^30^) with a sensitivity and specificity > 99% (see Supplementary Methods section 2 for details). On average the individual mites laid 9 (± 2.3) eggs on bean, 3.5 (±1.8) eggs on tomato during the first 24h and 2.9 (±2.7) eggs between 24h and 48h on tomato.

As shown in Supplementary Figure 2, fecundity on bean is strongly correlated with fecundity on tomato during the first 24 hours (p = 2.5 × 10^−05^), likely reflecting residual effects of prior feeding on bean. This relationship is much weaker for fecundity between 24 and 48 hours on tomato (Pearson R = 0.13, p = 0.07), indicating that host-specific performance becomes more apparent during the second interval. Although this correlation is not statistically significant, it may still capture differences in general fitness. We therefore included fecundity on bean as a covariate in subsequent fecundity analyses (see Section 2.5) to control for residual differences in overall fitness and to focus specifically on tomato performance.

### 2.3 Genotyping of the multiparental mapping population

Whole genome sequencing (2×150 bp paired-end) was performed for the 6 iso-female lines, and RNA-seq reads (polyA+, 2×150 bp paired-end) were collected from 190 single female mites from the mapping population. GATK’s best practices workflow^31^ was used for DNA genotyping of the parental founders and initial RNA genotyping for the mapping population, followed by the application of a HMM to identify the parental origin along the genome of each mapping population sample (Figure 1C). Of the ∼3.5 million raw genetic variants identified in the parental WGS data, 202,533 variants met the criteria for HMM analysis. These variants were homozygous in all parents, differed between at least two parents, and passed the additional filtering criteria outlined in Supplementary Methods sections 1 and Methods section 5.3.1.

Figure 2C shows the parental proportions in the mapping population along the genome, as defined by the HMM results. Although the parental proportions in the mapping population followed similar distributions across the chromosomes, except from some distortions in smaller regions, they were imbalanced (Figure 2C; Supplementary Table 1). The proportions of lines PL and UK were clearly elevated, with median proportions across the three chromosomes being 0.38 and 0.26, respectively. The ES parental line was barely represented in the mapping population, with a median proportion of 0.01 across the chromosomes. These proportions, inferred from the HMM-based parental origin, align with those calculated from the allele frequencies of unique parental marker SNPs in a pooled sample of the *quasi*-panmictic population after 3 generations (see also Villacis-Perez et al.^25^; Supplementary Table 1), which were also used as initial probabilities in the HMM (see Methods section 5.3.1).

**Figure 2.**
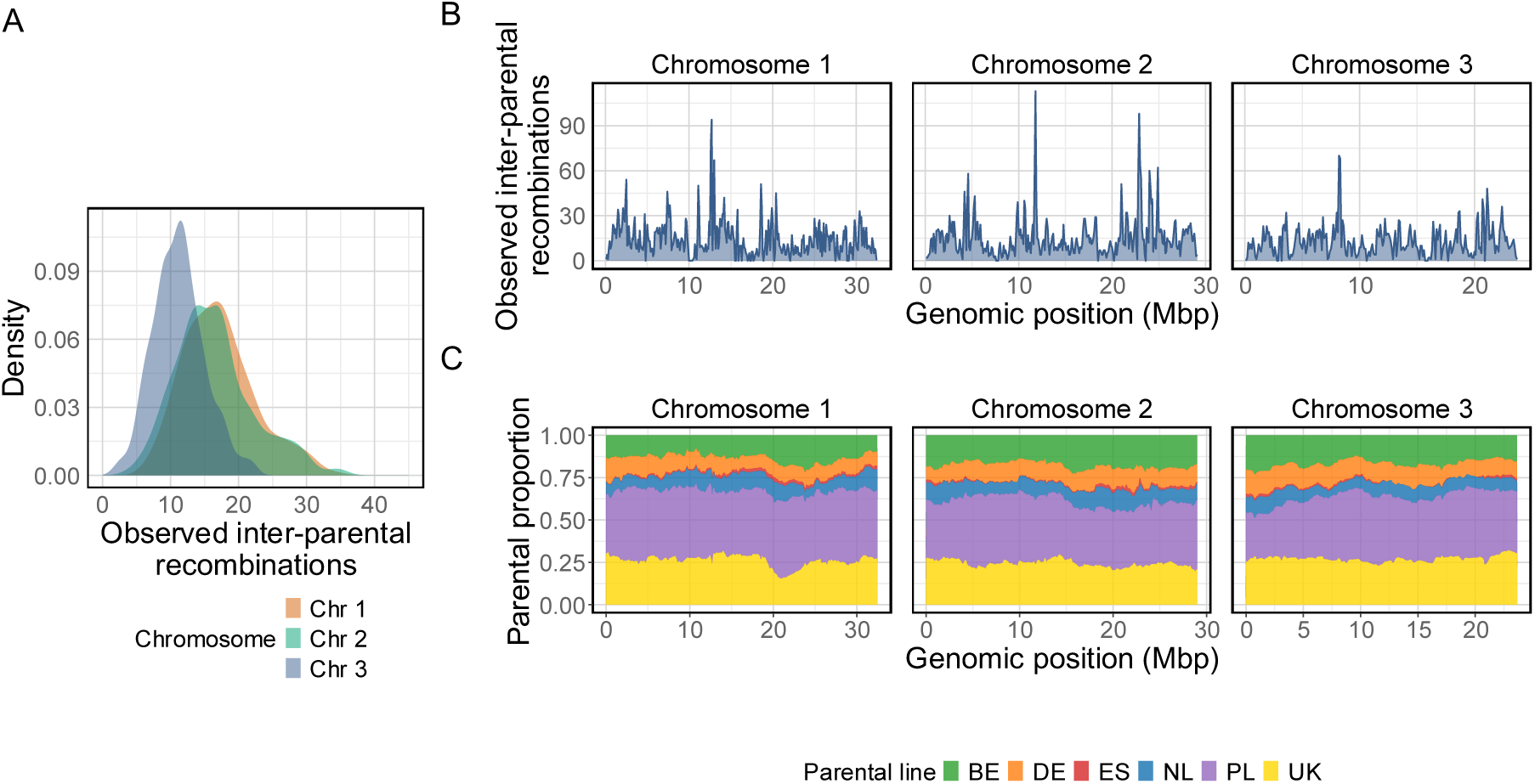
Distributions of observed inter-parental recombination events and parental proportions within the mapping population. A) Density plots of the observed inter-parental recombination events per chromosome and per sample, as determined by the parental shifts identified by the HMM. B) Distribution of observed inter-parental recombination along the 3 chromosomes, assessed with a sliding window (window size = 200 kb, step size = 100 kb). C) Distribution of parental proportions along the 3 chromosomes, based on the parental origin of all samples determined by the HMM in sliding windows (window size = 200 kb, step size = 50 kb). Colors represent the parental iso-female lines, as indicated in the legend.

Despite the moderate imbalance, we proceeded with the mapping population for developing and applying the eQTL methodology. Based on the HMM, shifts in parental origin along each individual chromosome can be identified, referred to as observed inter-parental recombinations. A total of 8,527 inter-parental recombination events were observed across all samples and chromosomes. The average number of inter-parental recombinations per sample was 44: 17 for chromosome 1, 16 for chromosome 2, and 11 for chromosome 3 (Figure 2A), proportional to the chromosomes’ relative lengths. Analysis of inter-parental recombinations in sliding windows (Figure 2B) did not reveal any large regions with very low recombination rates. By assuming panmixia, the average number of recombinations since the creation of the mapping population (including recombinations between alleles of the same parent) could be estimated at 1.7 recombinations per generation per chromosome (see Supplementary Methods section 5 for details), only slightly lower than the observed median number of 2 recombination events for the 3 *T. urticae* chromosomes reported previously^11^, possibly due to the inability to detect very small recombination events.

Subsequently, the mapping population’s genomes were reconstructed as a combination of parental haploblocks. DBGs were constructed for the founding parents, allowing for at most 2 parental alleles per gene. This was successful (see Methods section 5.3.2 for criteria) for on average 15,199 (± 106) out of 18,223 genes (∼84%) per parent. On average, 10% (UK) to 42% (PL) of all genes featured more than one assembled allele, reflecting varying levels of homozygosity, though it should be noted that this is a lower estimate given the need for genetic variation within the genes. Next, in order to complete missing gene-level information and to remove technical artefacts, mapping population samples were reconstructed as a function of parental haploblocks (see previous section and Methods section for more details). Genotyping of heterozygous parental haploblocks was successful for on average 81% (± 8%) haploblocks. For the remaining haploblocks we could not discriminate between parental alleles and these were hence collapsed into a single allele. Overall, we obtained an average of 883 ± 110 haploblocks per individual (diploid genome) in the mapping population, with a mean missing genotype fraction of 0.017 (± 0.005) per sample. Note that this number of haploblocks also encompasses recombination events that occurred in the iso-female lines prior to constructing the *quasi*-panmictic population.

The union of all haploblock borders was used to define a total of 5,708 genotype bins, i.e. genomic regions without any recombination events in the mapping population. The use of haploblocks indeed removed artefacts caused by the use of RNA-seq data for genotyping, as demonstrated in Figure 3. For loci where all genotyped samples have matching genotypes, both the haploblock-based and GATK approach yield a distribution consistent with Hardy-Weinberg equilibrium (Figure 3A). In contrast, GATK-based genotypes clearly deviate from this pattern when mismatches occur, with the deviation becoming more pronounced as the number of mismatches increases (Figure 3B-D). Thus, the haploblock-based strategy effectively eliminates RNA-based genotyping artefacts.

**Figure 3.**
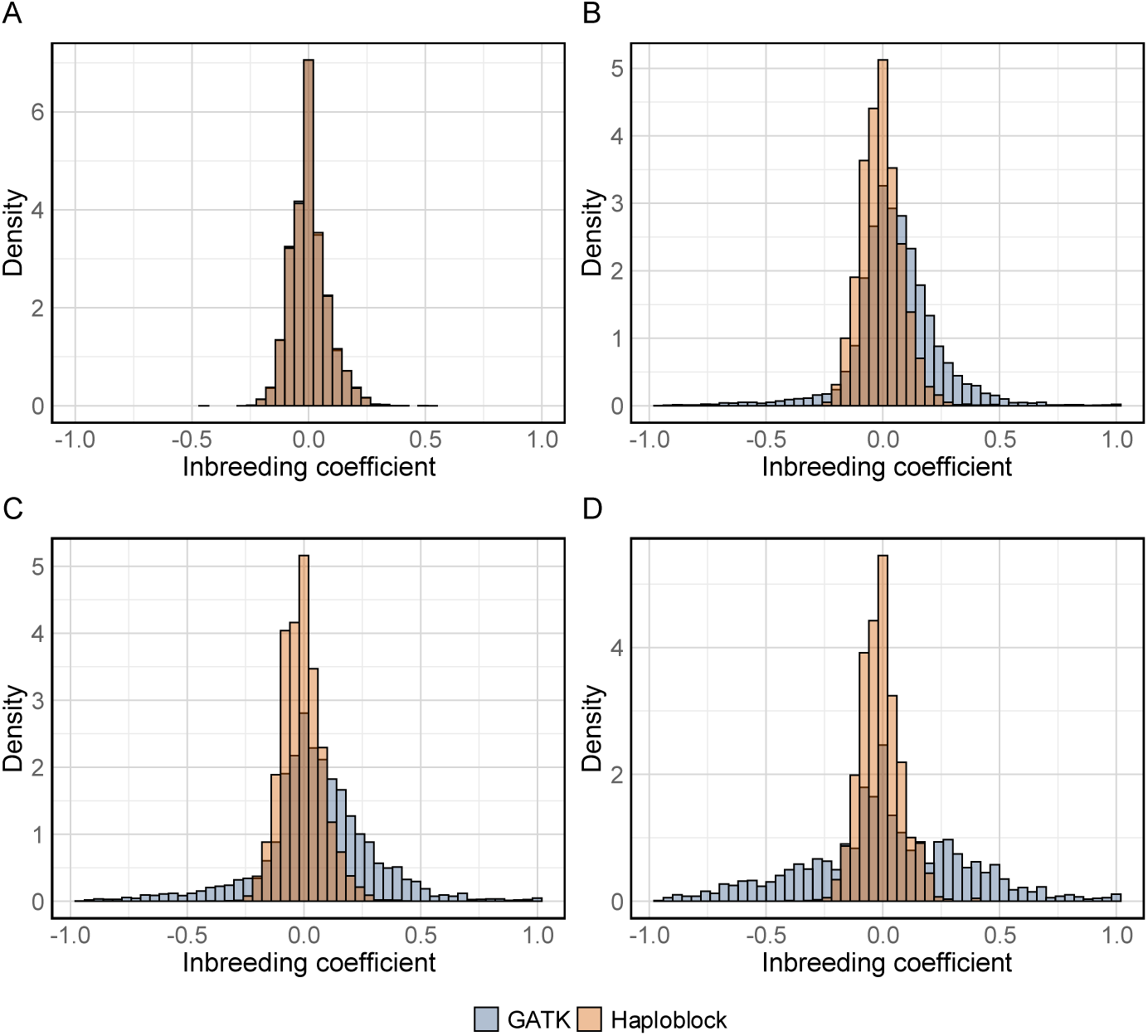
Histogram of inbreeding coefficients for subsets of bi-allelic SNPs comparing the haploblock-based genotypes (orange) and RNA-seq-based GATK genotypes (blue). Subplots show: A) matching SNPs (n = 44,258), B) SNPs with >5 mismatches (n=14,275), C) SNPs with >10 mismatches (n=6,370), and D) SNPs with >25 mismatches (n=2,217).

### 2.4 eQTL mapping

In a *quasi*-panmictic population, genetically determined gene expression may be determined by multiple *cis* and *trans* effects. We therefore screened for eQTLs using a regularized regression approach (LASSO, 25x repeated 4x cross-validation), which predicts gene expression levels by considering all parental genotype variables, filtered on a minimal minor allele fraction of 0.05 (n = 9,823), simultaneously. Average predictions from the test datasets were used to estimate the proportion of expression variance explained (R²) by the genotypes (Figure 4A), and only eQTLs for genes with an R² > 0.01 were further considered. Features selected in at least 65 of the 100 models, and with consistent effect directions in all 65 models, were classified as eQTLs for the corresponding gene. We repeated this procedure with permuted data, to confirm that gene expression could not be predicted using the latter (Figure 4A), and to provide an indication of the expected number of false positives.

**Figure 4.**
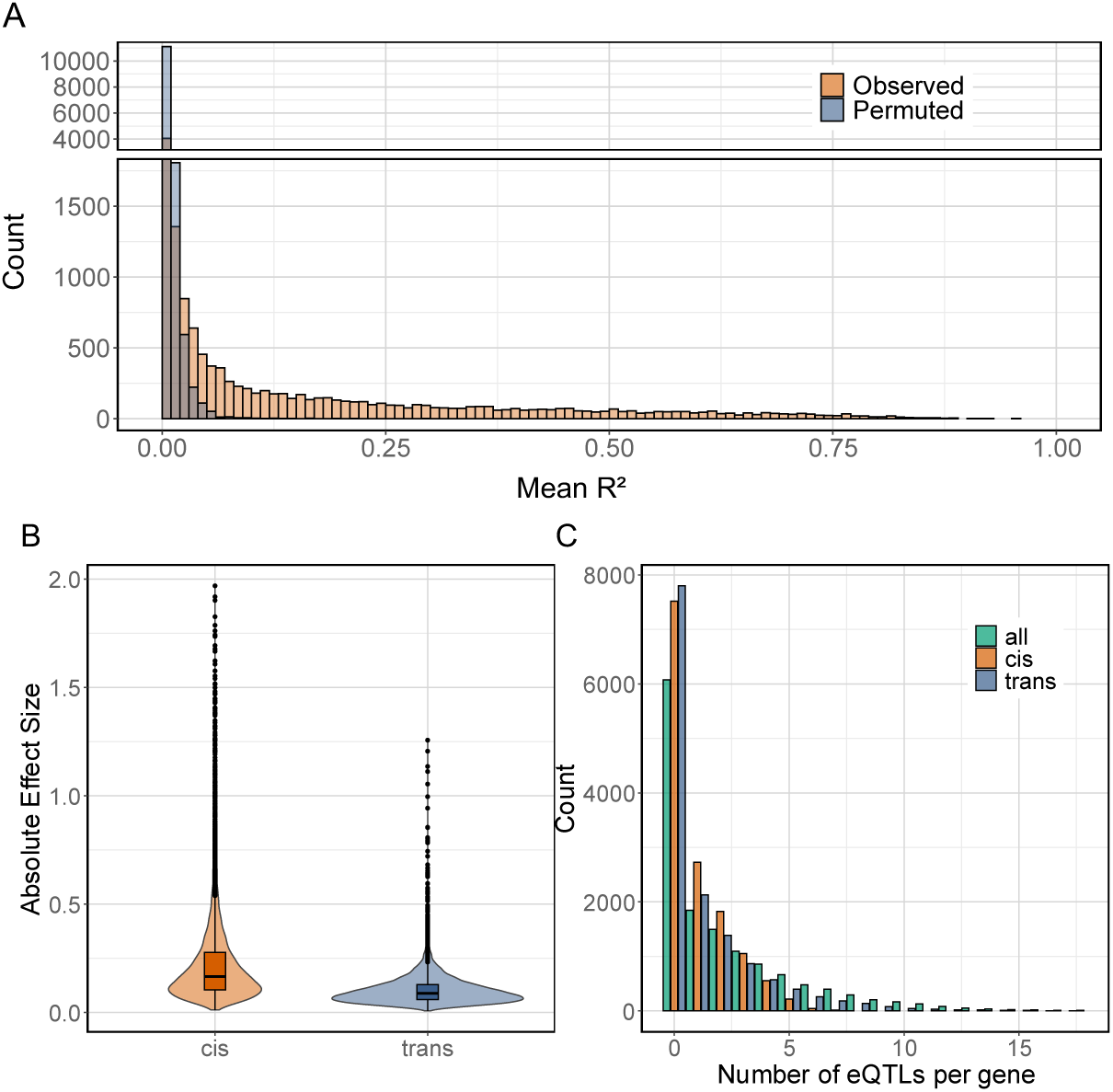
Characteristics of eQTLs in a *quasi*-panmictic, multiparental mapping population using regularized linear regression. A) Mean R² of all genes derived from linear regression models predicting gene expression based on the genetically determined gene expression (LASSO predictions) for the actual, observed data (orange) and permuted data (blue). B) Mean absolute effect sizes (beta values from LASSO models) for *cis*- and *trans*-eQTLs. C) The total number of eQTLs (green; *cis* + *trans*), *cis*-eQTLs (orange) and *trans*-eQTLs (blue) per gene.

In total, we identified 33,029 eQTLs of which 31,236 were linked to target genes (n = 7,869) located on the 3 chromosomes (Figure 5A; we excluded those located on small, unplaced scaffolds). It should be noted that our haploblock strategy entails that eQTL variants shared between parents will be detected on a per parent basis, and hence be included multiple times. For the permuted data, where the exact same model fitting and eQTL selection procedure was applied, 290 false positive eQTL bins were detected (see Methods section 5.4.2 for details), implying that approximately 0.9% of the eQTLs identified in the observed data are expected to be false positive. Of the identified eQTLs, 13,174 (42%) were classified as *cis-*eQTLs, which we defined as those for which the midpoint of the bin of maximal association was within 1 Mb of the target gene, while 18,062 (58%) were otherwise classified as *trans*-eQTLs. Of the target genes, 1,724 (22%) were regulated solely by *cis*-eQTLs, 1,442 (18%) solely by *trans*-eQTLs, and, 4,703, (60%) by both *cis*- and *trans*-eQTLs, with 2,371 (30%) regulated by multiple *cis* and multiple *trans* effects. The number of eQTLs per gene are depicted in Figure 4C. On average, *cis*-eQTLs had larger absolute effect sizes than *trans*-eQTLs (0.22 vs. 0.10) (Figure 4B), consistent with previous findings^4,11,32^.

**Figure 5.**
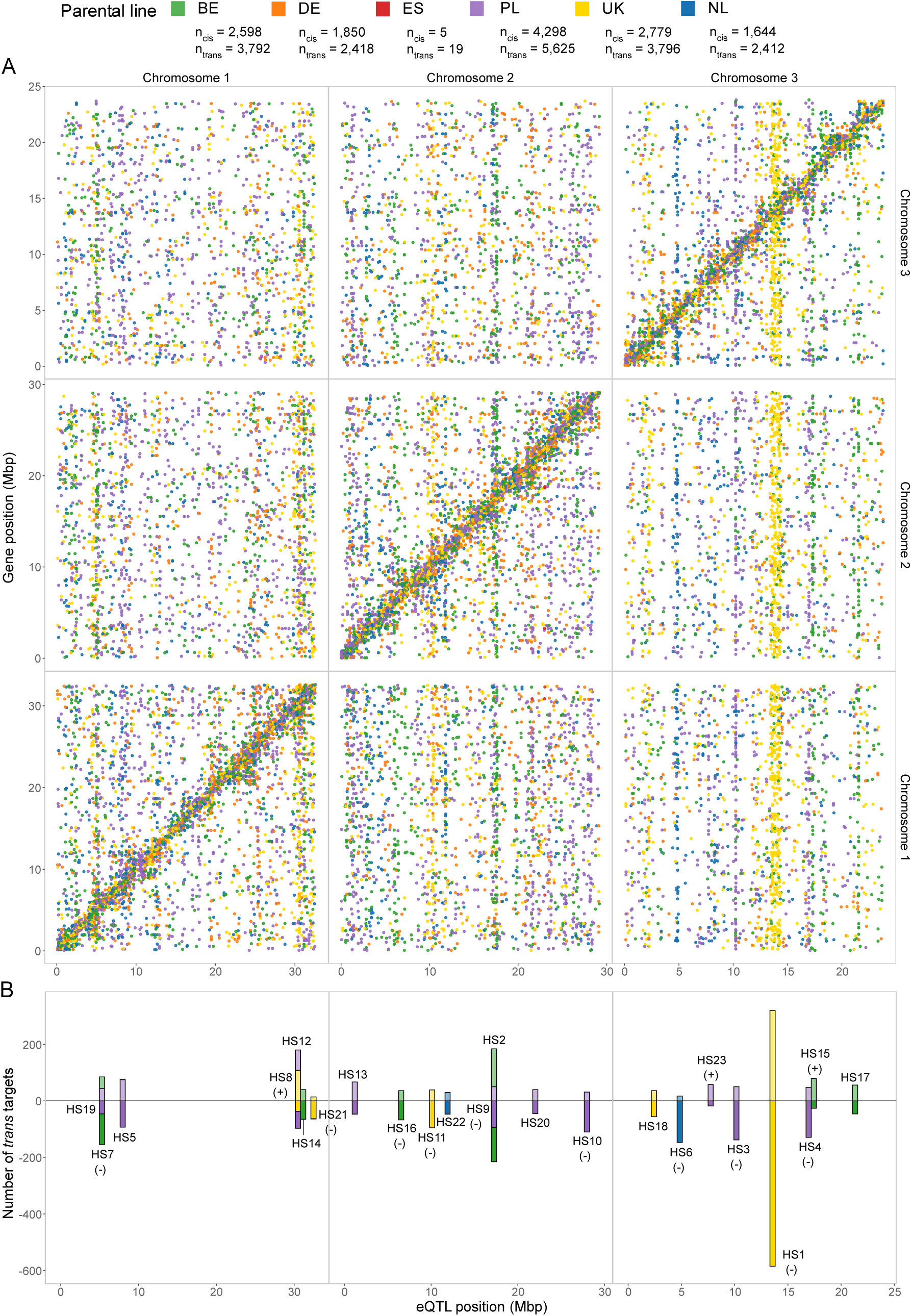
eQTLs in a *quasi*-panmictic, multiparental mapping population using regularized linear regression. A) Scatter plot showing eQTLs and the corresponding position of the eQTL bin (x-axis) and target gene (y-axis). Colors indicate the parental line of the eQTL as shown in the legend. The legend also shows the total number of *cis*- and *trans*-eQTLs detected per parental line. B) Hotspot regions regulating more than 75 genes in *trans*. The number of upregulated genes is shown along the positive y-axis (light shade), and the number of downregulated genes is shown along the negative y-axis (dark shade). Each hotspot is labelled with its ID, positioned according to the majority regulation (up or down), and includes a + or – sign if a significant bias (ꭕ², Bonferroni-adjusted p-value < 0.05) towards upregulation or downregulation is observed for the *trans* effects.

For context, results from the single-variant (MatrixEQTL^33^) method are shown in Supplementary Figure 3. Using an FDR threshold of 0.01, MatrixEQTL identified 23,178 significant associations across 8,273 genes on the three chromosomes. The proportion of *cis*- and *trans*-eQTLs differed significantly from the LASSO results (Fisher exact test, p < 2.2×10^−16^), with 54% *cis* and 46% *trans* associations. Of the detected eQTLs, 70% overlapped between the two approaches, and shared signals were particularly enriched for *cis* effects (∼66%). eQTLs unique to MatrixEQTL tended to have smaller effects/significantly larger p-values (permutation test, p < 1×10^−07^), whereas genes identified only by LASSO were typically influenced by multiple loci (permutation test, p < 1×10^−07^). Because the two approaches differ fundamentally in their statistical frameworks – MatrixEQTL uses hypothesis testing with FDR control, whereas LASSO selects predictors via cross-validation − their results should not be interpreted as directly comparable. Nonetheless, the general trends suggest that MatrixEQTL is more sensitive to single small-effect loci, whereas LASSO captures genes affected by multiple regulatory variants.

Based on the LASSO eQTL results, we identified *trans* regulatory hotspot regions by quantifying *trans*-eQTLs per parental line in non-overlapping 300 kb windows. We obtained 23 distinct hotspot regions, defined as genomic regions regulating more than 75 genes in *trans* (Supplementary Table 2), and additionally evaluated their impact on detoxification gene expression.

The most striking hotspot was HS1 (UK, chromosome 3, ∼13.8 Mb, 905 target genes, 81 detoxification genes (see Supplementary Methods 4 for the definition of detoxification genes), Figure 5B, Supplementary Table 2-3) covering a relatively broad region and having the largest number of *trans* associations. For the targets of this hotspot we noted general expression downregulation when the UK allele (a1) was present (ꭕ² p-value = 9.39×10^−19^). The hotspot featured multiple significantly enriched GO terms (FDR < 0.05) among its target genes (Supplementary Table 4), including GO:0036094 (small molecule binding), GO:0031409 (pigment binding), GO:0004869 (cysteine-type endopeptidase inhibitor activity), GO:0016702 (oxidoreductase activity), GO:0004197 (cysteine-type endopeptidase activity) GO:0008234 (cysteine-type peptidase activity), GO:0016887 (ATPase activity), and GO:0043401 (steroid hormone mediated signaling pathway), all related to the detoxification of plant metabolites and digestion. The terms GO:0036094 and GO:0031409 correspond to the detoxification genes of the lipocalin family. While the HS1 region contains many genes, candidates to underlie the *trans* effects include *tetur11g01Sc0* (a canonical HR96 nuclear receptor), which harbors a serine-to-proline substitution and a proline-to-serine substitution and *tetur11g00260*, a Krüppel-like zinc finger with a proline-to-leucine substitution. In both cases, the substitution is specific to the UK eQTL allele. In addition, this region features a large and complex cluster of copy number variable putative limkain-b1 genes, a poorly studied family in arthropods that may play a role in gene regulation, as indicated by the assigned GO terms. Lastly, this hotspot region also had significant *cis*-eQTL effects on *tetur11g01430*, which encodes a negative elongation factor E, and *tetur11g00180*, a suppressor of Hairless. The expression levels of these *cis*-regulated genes correlate significantly with those of the *trans*-regulated targets, making them strong candidates for assessing whether the *trans* effects are mediated through *cis*-regulation of these genes.

A second notable hotspot is HS6 (NL, chromosome 3, ∼4.9 Mb, 164 genes, 16 detoxification genes, Figure 5B, Supplementary Table 2-3) which also exhibited a significant bias in direction (expression downregulation) of the *trans* effects when the NL allele was present (ꭕ² p-value = 3.27×10^−24^), and several GO terms were enriched among its target genes (Supplementary Table 4): GO:0008234 (cysteine-type peptidase activity), GO:0004553 (hydrolase activity), GO:0043169 (cation binding), GO:0016705 (oxidoreductase activity), GO:0005319 (lipid transporter activity), GO:0006508 (proteolysis), and GO:0005975 (carbohydrate metabolic process). Within this eQTL hotspot region, we identified the initiation factor eIF-4 gamma (*tetur02g11cc0*), which exhibited two significant amino acid changes specific to the NL parental line (serine-to-isoleucine and glutamate-to-glycine). In addition, this region contains several orthologs of Argonaute-1 (*tetur02g10580, tetur02g10570, tetur02g10560*), genes that are key in RNA silencing in *Drosophila* and multiple other insect species^34^. However, CNV within these genes complicates the interpretation of coding changes.

Given our interest in host plant use, we further identified *trans*-eQTLs featuring major fractions of detoxification genes among their targets. A notable region was located around 12.5 Mb on chromosome 1, where both the DE (21 genes, 11 detoxification genes) and NL (71 genes, 16 detoxification genes) parents exhibited high fractions of detoxification genes among their target genes (52% and 23%, respectively; Supplementary Table 5). This locus contains *tetur06g04270*, an HR96 nuclear receptor with a ligand-binding domain but lacking the conserved DNA-binding domain, and was previously identified as a *trans*-regulatory hotspot for detoxification genes^11^. In our study, 10 of the 11 detoxification targets in this eQTL region for the DE parental line overlapped with *trans* detoxification targets identified in the earlier study, including the P450s: *CYP392D2, CYP392D3, CYP392D8, CYP3S2DSP*, and *CYP392D5p*. All overlapping genes were upregulated when the DE allele was present, with a mean beta effect size of 0.23, more than twice the average effect size for all *trans* associations (0.11), further supporting the particularly large effect sizes associated with this hotspot. Similarly, in the NL parental line, 16 detoxification genes were targeted by the same region, 12 of which overlapped with previously identified targets of the HR96 receptor, including the shared P450s observed in the DE parental line and some additional P450s (*CYP392A12* and *CYP392A11*). Based on the DBG assemblies of the parental alleles and the corresponding Illumina WGS reads, we identified a T-to-A transversion that leads to the same W309R (tryptophan-to-arginine change) previously observed in the multi-pesticide resistant strain used in the Ji et. al.^11^ study in parental line NL, but not in line DE.

### 2.5 *Trans*-eQTLs underlying spider mite fitness on a new host plant

Next, we evaluated whether identified *cis*- and *trans*-eQTLs contributed to the phenotype under study, i.e. fitness on a new challenging host (tomato), quantified as fecundity. Per gene, genetically determined expression was used as predicted by the LASSO model (see 2.4). Linear regression was then applied to identify genes for which fecundity on tomato was associated with genetically determined gene expression upon adjustment for fecundity on bean. Consistent with this design, haplotype GRM–based heritability was near zero for fecundity on bean during the first 24h and for fecundity on tomato during the first 24h, but was moderate for fecundity on tomato between 24 and 48h (h² = 0.16), indicating that genetic effects on fecundity become detectable only after extended exposure to the tomato host. Although bean fecundity itself shows little additive genetic variance, it still captures residual differences in general fitness that can influence performance after transfer (see Section 2.2). This approach therefore isolates tomato-specific variation in fecundity, as expression was measured on tomato, where transcriptional responses occur rapidly, while fecundity adjusts more slowly and can remain affected by prior feeding on bean.

To assess patterns of phenotype association for *cis*- and *trans*-regulated genes, we selected genes with substantial genetically determined gene expression (R² > 10%) and created subsets of genes categorized as predominantly *cis* regulated (n = 2,237), predominantly *trans* regulated (n = 368) or regulated by a mix of *cis* and *trans* mechanisms (n = 2,546) (see Methods section 5.5 for details).

Fitness on tomato was predominantly associated with *trans*-eQTL effects, with a substantial number of genes for which genetically determined expression explained fecundity on tomato more than expected by chance (Figure 6A). In contrast, for genes predominantly regulated in *cis* (Figure 6C), or by a mixture of effects (Figure 6B), genetically determined expression showed far less association. As a negative control, we performed a comparable analysis with genes with poor predictive value of gene expression (R² < 1%, n = 4,063), and found no deviation from what was randomly expected (Figure 6D).

**Figure 6.**
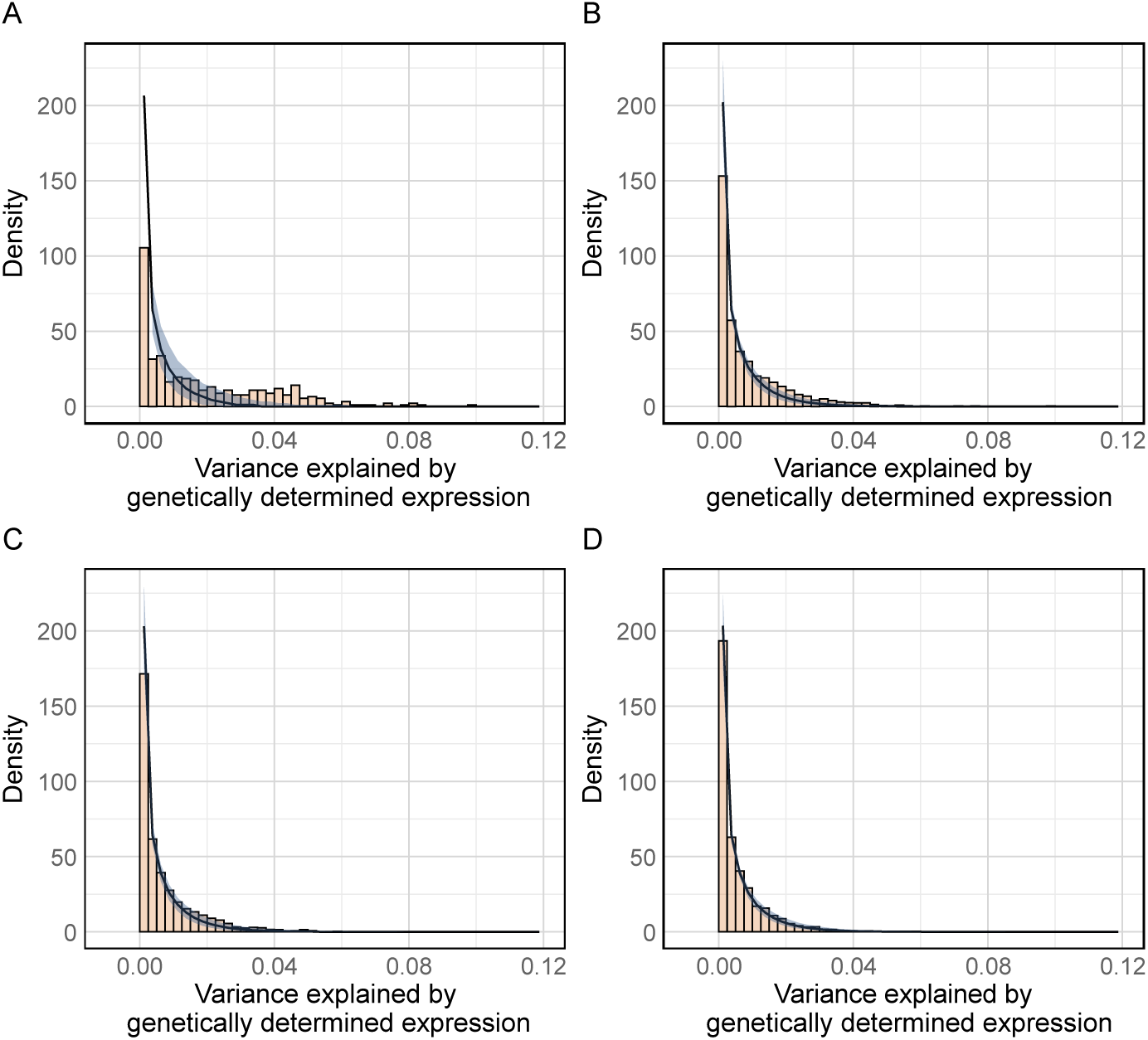
Histograms showing the variance in fecundity on the challenging host tomato explained by genetically determined expression. The observed data are displayed as histograms, with the black density line representing the median trend under the null hypothesis and the blue shaded area indicating the 95% confidence interval. Subgroups are defined based on gene regulation: A) Predominantly *trans* regulated genes (n = 368), B) Genes with mixed regulation (n = 2,546), C) Predominantly *cis* regulated genes (n = 2,237), D) Genes with limited genetically determined gene expression (n = 4,063).

Among the predominantly *trans*-regulated genes, we identified 89 genes whose genetically determined expression significantly predicted fecundity on tomato (FDR < 0.05) (Supplementary Table 6). In contrast, no significant associations were observed for predominantly *cis*-regulated genes after multiple testing correction, and only 3 genes were significant among the mixed-regulated genes. Nevertheless, Figure 6 suggests that increased power would reveal more significant results for at least some of these smaller effects for non-predominantly *trans*-regulated genes. A similar analysis for the initial time interval on the new host (24h after transfer) revealed minimal genetic regulation of fecundity at this stage (Supplementary Figure 4), likely due to the still evolving defensive response of tomato, remaining effects of feeding on bean as well as the use of RNA profiles at 48h on tomato. The results were also highly consistent with those from the analysis without correction for fecundity on bean (Supplementary Table 7), with 84 of the 89 significant genes overlapping between the two approaches.

For the 23 hotspot regions (Results section 2.4), we assessed whether fecundity-predicting genes were enriched among the *trans* targets of each hotspot, using the predominantly *trans*-regulated genes (n = 368) as background (Supplementary Table 8). Six hotspots (HS1, HS2, HS15, HS17, HS19 and HS23) showed significant enrichment of phenotype-associated *trans* targets (Fisher exact test; Bonferroni adjusted p < 0.05). The most striking result was for HS1, also described in Results section 2.4, which showed a major enrichment for *trans* targets associated with fitness on tomato (p = 1.92×10^−31^; odds ratio (OR) = 30). Presence of the UK allele (a1) was associated with expression downregulation for 71 (90%) of these fitness associated *trans* targets, an enrichment (Fisher exact test OR = 4.97, p = 2.48 × 10^−04^) compared to all predominantly *trans* regulated targets for this hotspot. Moreover, for 70 (89%) of the fitness associated *trans* targets of this hotspot, the genetically determined expression was negatively correlated with fecundity on tomato. These results suggest that presence of the a1 allele leads to higher fitness and fecundity, with a lower general stress response (hotspot target gene downregulation) as indirect consequence.

We additionally used a traditional QTL mapping approach to verify whether the genotype at each phenotype-associated hotspot region was significantly associated with fecundity on tomato at 48h (Supplementary Table 9). This was the case for the genotypes at hotspot regions HS1 (p = 0.0008), HS2 (p = 0.01), HS15 (p = 0.009) and HS17 (p = 0.01), which all ranked among the top 2.5% of results in the QTL analysis. Note that these loci were not significantly associated with fecundity on bean (all p-values ≥ 0.33) or tomato during the first 24h (all p-values ≥ 0.21). Combined (assuming additive effects and adjusted for bean fecundity), these loci explained about 11.1% of total variation in fecundity on tomato. HS19 (p = 0.10) and HS23 (p = 0.15) did not show a significant association. Importantly, none of these hotspot loci were significant using a genome-wide QTL strategy due to multiple testing correction. These results demonstrate that the strategy outlined in this study can successfully identify loci associated with fecundity.

## 3. Discussion

In this study, we present a cost-efficient strategy for eQTL detection that addresses several common challenges in experimental eQTL studies. Using a multiparental population and experimentally simple and cost-efficient design in a proof-of-principle *T. urticae* study we were able to identify multiple *trans*-eQTL hotspots. Moreover, some of these were also significant QTLs for fecundity on a major crop host plant. Even for *T. urticae*, a well-studied organism given its agricultural relevance and straightforward manipulation, direct screening for (e)QTLs associated with an agriculturally important phenotype is unique.

We used heterozygous individuals to construct a multiparental mapping population. As expression of a gene may be affected by multiple eQTLs, relying on a genetically diverse population was suggested to complicate *trans-*eQTL mapping^5^. Therefore, we used a regularized regression strategy (LASSO) to evaluate the combined impact of multiple smaller eQTLs effects. Far more *trans*-eQTL effects were observed than when using a single-variant regression strategy, which predominantly identified *cis*-eQTLs. These single-variant results contradict previous studies reporting more *trans* than *cis* effects in the two-spotted spider mite^11,19^. Nevertheless, it should be noted that differences in sample size (power), practical *cis/trans* definition, and technical biases caused by e.g. alignment issues all complicate an exact comparison between studies.

In this study, power issues in detecting *trans*-eQTLs were addressed by reducing multiple testing through a haploblock strategy. This haploblock strategy was also required to eliminate artefacts (e.g. alignment errors) from the raw RNA-seq based genotyping results, which may otherwise lead to false positive eQTLs, a problem also arising in DNA sequencing based strategies^35^. While the concept of haploblocks is not new and has previously been applied for eQTL mapping in a biparental *T. urticae* population by Ji et al.^11^, as well as in QTL mapping studies using multiparental populations – including heterogeneous stock^20^ and diversity outbred^21,22^ mice populations and multiparent advanced generation intercross (MAGIC) populations in Arabidopsis^23^ and maize^24^ – our methodology differs from most of these approaches in not requiring inbred parental founders or complete haplotype information, and in relying on RNA-seq for haploblock-based genotyping of the mapping population.

Our experimental design for eQTL mapping provides significant advantages over traditional methods. In particular, it markedly reduces the laboratory workload compared to eQTL mapping designs that require hundreds of individual crosses (e.g. Ji et. al^11^), and it does not require the time-consuming step of generating highly inbred lines. The sole condition is that a *quasi-*panmictic population can be created from founders with sufficient genetic variability. In this study, iso-female lines were used as founders because individual-level genotyping is not feasible given the minute size of spider mites. These lines are partially heterozygous but contain at most two different alleles, enabling population-level sequencing that approximates heterozygous individuals without phasing information. The fraction of remaining heterozygosity fluctuated for these iso-female lines, reflecting that our methodology can start from any limited set of heterozygous or (partially) homozygous/inbred individuals. However, the approach would be considerably simplified if fully phased haplotypes of individual founders were available, as this would allow direct inference of haplotype pairs in a single HMM step. Additionally, our eQTL mapping strategy predominantly relies on RNA-seq, thereby drastically reducing costs and experimental complexity than when DNA-based genotyping is additionally required. Even with decreasing sequencing costs, available budgets can hence be more efficiently used, e.g. to increase mapping population sizes to enhance statistical power, or for collection of richer phenotypic data.

Nevertheless, several considerations should be made with applying the strategy introduced here. First, it should be possible to create a *quasi-*panmictic population, i.e. without any genetically distinct subpopulations. In our case, this entailed two initial crosses between parents. Even then there was imbalance in the contribution of each parental genotype to the final mapping population, which could be caused by e.g. fitness differences or specific incompatibilities between parents. This imbalance will negatively affect power to detect the allelic impact of the less represented parents and should hence be minimized. This can be easily monitored by WGS of a single pooled sample of the population. Nevertheless, as long as the effect is moderate, it will not directly affect the accuracy of identified eQTLs. Other considerations largely depend on the research question and experimental practicalities. For example, screens focusing on the identification of larger effects can incorporate more parents and/or smaller sample sizes, while the comprehensive characterization of all genetics effects may need to maximize power by using fewer parents and larger sample sizes. Additionally, recombination rate, genetic variability of the parents and the number of generations used to create the mapping population play a major role to obtain optimal resolution. For larger bins, e.g. HS1 in our study, it is harder to pinpoint the underlying gene, and even the presence of two linked genes affecting phenotype cannot be excluded. A higher recombination rate and number of generations will increase the number of recombination events and hence the mapping resolution, but also lead to stricter multiple testing correction and requires more genetic variation between parents to resolve smaller haploblocks. Attention should also be paid to false positive eQTLs: these will be overfit to the gene’s expression, and may thus lead to an association with phenotype independent of genetics. To mitigate overfitting in eQTL discovery using the LASSO methodology, we employed cross-validation and stability selection, and used permuted data to obtain an estimate of the number of associations expected due to overfitting.

Using this novel methodology, we focused on identifying *trans*-regulatory hotspots, with a special interest in hotspots affecting the expression of detoxification genes. These genes are critical for the adaptability of *T. urticae* to new host plants and the development of pesticide resistance^15,16,27^. Despite their significance, the genetic mechanisms regulating these detoxification genes remain only partially understood. Interestingly, we detected major *trans*-eQTL effects near an HR96 nuclear receptor with a ligand-binding domain (but lacking a DNA binding domain) for a DE and NL parental allele that regulated mostly detoxification genes, primarily P450s. Underscoring the robustness of our methodology, Ji et al.^11^ identified the same gene as a major *trans*-eQTL hotspot regulating detoxification in an acaricide resistant line, with substantial overlap of targets. Moreover, in parental line NL a mutation causing the same W309R (tryptophan-to-arginine) substitution as identified by Ji et al.^11^ was observed.

While this study identified numerous genomic regions as *trans*-regulatory hotspots, the individual candidate genes affecting transcription remain to be experimentally validated. Follow-up experiments using e.g. RNAi or CRISPR/Cas9^36^ are hence essential to validate candidate genes and uncover the underlying molecular biology. From a mechanistic perspective, it should be noted that the *trans* targets per hotspot are not necessarily directly transcriptionally regulated through the eQTL gene, as indirect effects may occur when the underlying genetic variant has a major impact on fitness on the challenging host. Nevertheless, even indirect effects are valuable for the here introduced eQTL strategy, since our overarching goal was exactly to identify such genetic variants.

## 4. Conclusion

In summary, we developed an RNA-seq driven strategy for genotyping individuals of a multiparental population when WGS data exists for the non- or partially-inbred parents. This provides a cost-efficient option for eQTL scanning and enables the identification of genotypes and *trans*-eQTL hotspots associated with specific phenotypes. Our study underscores the importance of investigating intra-specific genetic variation in gene expression on phenotypes, in our test case fitness of a generalist mite herbivore after a shift to a challenging host plant.

## 5. Methods

### 5.1 Mite population and DNA sequencing

The characteristics of the mapping population used in this study were previously summarized by Villacis et al.^25^. Six iso-female lines of *T. urticae* were derived from field strains, collected from different locations and hosts in Europe, through expansion of a cross between a single female and her son. Genomic DNA (gDNA) was retrieved from a sample of 400-800 mites per parental line for sequencing and variant calling (see Supplementary Methods section 1).

The 6 lines were used to create a *quasi*-panmictic population, hereafter referred to as the mapping population, following the crossing scheme depicted in Villacis-Perez et al^25^. Briefly, the six lines were divided into three crossing blocks, each of which was comprised by ten virgin females from one line, ten virgin females from a second line, ten males from a third line and ten males from a fourth line. Then, 20 virgin females from the resulting F1 generation from each block were crossed with 20 males of the remaining two lines (i.e. fifth and sixth line). The F2 offspring obtained from these crosses, which were hence heterozygous throughout their genome, were transferred together to multiple bean plants to expand. The mapping population was maintained on potted bean plants (*Phaseolus vulgaris* L. cv. Prelude), in a climatically controlled incubator at 25°C (±0.5°C), 60% relative humidity, and 16:8 light:dark photoperiod. A pooled mapping population sample (same procedures as for parental lines) was sequenced to estimate the contribution of each parental line.

### 5.2 Single female sampling, RNA sequencing and phenotyping

Three hundred virgin females (teleiochrysalid stage) of the mapping population were isolated with some males, to ensure fertilization, after at least 8 generations. They were allowed to grow into 5-day-old females on detached bean leaves placed on wet cotton before transferring each female to a separate bean leaf disk (*Phaseolus vulgaris* L. cv. Prelude, 9 cm²). After 24h, each surviving female was transferred to a tomato leaf disk (*Solanum lycopersum* cv. Moneymaker, 9 cm²). The females were left on tomato leaf disks for 48h before freezing each surviving female separately (−80 °C). RNA extraction, sequencing, alignment and variant calling are further elaborated in Supplementary Methods. To assess the fitness of female mites on each host, we automatically counted the eggs per female laid every 24h (see Supplementary Methods section 2).

### 5.3 Decomposition of mapping population genomes into parental haploblocks

#### 5.3.1 HMM to identify samples’ parents along the genome

Based on the WGS-derived genotypes of the parental founders and the RNA-seq based genotype probabilities of the samples, we identified the samples’ parental origin across the genome using a HMM. In this model, each hidden state corresponds to a specific combination of founding strains. Since we started from 6 different iso-female lines, there are 21 possible combinations or hidden states.

Each node in the HMM corresponds to a high-quality bi-allelic variant that met the filter criteria specified in Supplementary Methods section 1, and that is located in putative single-copy regions (described in Supplementary Methods section 3.2). Furthermore, variants were only included if they segregated, had a WGS coverage ≥ 30 in the parental lines, and were homozygous in each parental line. Additionally, upon application of the HMM on the samples’ RNA based genotypes, a genetic variant was only considered by the HMM if the sample’s RNA coverage for that variant was ≥ 15 and if there was only a single most likely genotype.

A HMM is specified by: 1) transition probabilities, 2) initial probabilities and 3) emission probabilities. The transition probabilities, i.e. the probabilities of changing from one state to another, were estimated per chromosome using the median recombination frequency reported previously^11^. Based on the experimental design, and accounting for cases where recombination does not lead to a change in the HMM state (see also Supplementary Methods section 5), we estimated the number of state shifts for chromosome 1 to be 20. For chromosomes 2 and 3, the estimated shifts were set to 18 and 16, respectively, accounting for their lengths relative to chromosome 1. We did not consider double parental shifts between nodes (i.e. transitions between states that do not share a common founder and would require two recombination events between the nodes). Although biologically possible, it is highly unlikely given the recombination fraction and the SNP/node density, hence the transition probabilities of these events were set to 0. Initial probabilities for each state corresponded to the proportion of each parental line in a sequenced pooled sample of the mapping population (see Methods sections 5.1 and Supplementary Methods section 1). The emission probabilities, i.e. the observation likelihoods indicating the probability of an observation being generated from a certain state, were based on the sample’s probabilities (according to GATK) for the genotypes corresponding to those states. Note that for the HMM, only SNPs homozygous in each founder were considered, implying that there is only a single genotype possible per state. To ensure robustness against technical artefacts leading to incorrect GATK genotype probabilities (virtually) equal to zero, which would have major impact on the HMM, minor modifications were applied to these probabilities before use as emission probabilities: (i) a minimal genotype probability of 0.025 was used for all genotypes, (ii) a genotyping error rate of 5% was imposed by multiplying the emission probabilities by “(1 – genotyping error rate)”.

Once the initial, transition, and emission probability matrices were constructed, we implemented the Viterbi algorithm to identify, per sample, the parental composition across the genome as the most probable path through the hidden states. To correct for very short segments with likely spurious states, we altered the state of segments shorter than 1000 bp located in between two larger segments with an identical state to the latter state.

#### 5.3.2 Reconstruction of parental alleles

For each founder and annotated gene, we extracted WGS reads aligned to the corresponding genomic region, filtering for a minimum mapping quality of 60 and excluding duplicates. These filtered reads were then used as input for the bcalm2 tool^37^, which constructs de Bruijn graphs (DBGs) from sequencing data. We set the *k*-mer size (the length of the nodes of the DBG) to 19 and the abundance-min parameter to 1 to retain all *k*-mers, even those occurring only once. The output of the bcalm2 tool (the set of unitigs), was then − together with the sequencing reads − processed by a custom version of the detox^38^ tool. This tool performs graph cleaning and repeat resolution. The graph cleaning process enhances the quality and accuracy of the DBG by removing errors and artefacts, such as sequencing errors or low-quality reads. This includes removing tips and bubbles. In the repeat resolution step, we also aimed to correctly assemble repetitive sequences using coverage and paired-end information. After running the detox tool, we obtained the final DBG sequences, which we further refer to as alleles, for each gene and founder.

For single-copy genes with sufficient linked genetic variation and without recombination events, typically one allele (if the iso-female line was homozygous at that locus) or two parallel alleles (if the line was heterozygous at that locus) were constructed. These alleles hence represent the possible alleles for the selected gene in a particular iso-female line. However, when recombination events occurred in heterozygous regions, or when genetic variation was limited (no reads spanning over two adjacent SNPs, obscuring linkage between both), the DBG strategy may not reconstruct alleles spanning the entire gene region, resulting in multiple allele blocks. In such cases, all possible alleles were constructed by combining these allele blocks in all possible manners, yet only considering the gene’s coding region.

Genes were excluded from allele reconstruction and subsequent steps under the following conditions: when no alleles were assembled in the gene region; when more than two parallel alleles were assembled in the gene region (e.g. due to multiple copies or alignment errors); when the number of possible alleles (based on all allele block combinations) was ≥ 64 (indicating a highly fragmented and likely spurious assembly); or when the missing value fraction (of total bases covered by the gene region) was ≥ 0.25 (due to large deletions or CNV).

#### 5.3.3 Aligning the mapping samples’ RNA-seq reads to parental alleles

For each expressed gene and mapping sample originating from different parental lines, we aligned the RNA reads with the assembled parental alleles. Using the HMM assigned parental origin of each sample (see Methods section 5.3.1), we identified the candidate alleles for each gene per sample (see Methods section 5.3.2). We then relied on the reads covering all SNP positions that differed between the candidate alleles. For each possible combination of candidate parental alleles at a given gene for a particular sample, we fitted a weighted linear model if at least three SNPs were sufficiently covered (≥ 7 reads). In this model, the independent variables indicate the presence (1) or absence (0) of the variant SNP allele in the parental alleles, and the dependent variable the allele fraction of the variant SNP allele based on the RNA-seq data, with the square root of the total counts at each SNP used as weights. By fitting this weighted linear model, we calculated a total R² value for each combination of parental alleles, estimating how well we could predict the allele fraction of a sample across SNPs given the presence of certain parental alleles. Additionally, by comparing full models (both parental alleles) with partial models (only one parental allele), we calculated partial R² values to measure how much variance was explained by each individual parental allele.

As described in Methods section 5.3.2, often more than 2 candidate alleles per gene per parental line were assembled. As each parent (derived from a single female) has a maximum of two alleles (single-copy genes), we first identified the two most likely alleles per gene per parent. Therefore, per gene, for each parent and each selected individual originating from a cross between the parent under study and any other parent, we first identified the parental allele (under study) leading to the highest total R² value (as explained in the previous paragraph). Subsequently, across all thus selected individuals for that parent, we identified the two alleles most often leading to the highest total R² values. This was performed for all parental lines, leading to (at most) two alleles remaining per parent. Then, we filtered the weighted linear model results obtained for all possible alleles per parent to those remaining alleles. For each sample, this leads to at most two partial R²-values (one per allele) per contributing parent.

Whereas the allele with the highest partial R²-value could be considered most likely to be correct for that parent, we continued working with a measure of probability per allele, obtained by dividing each partial R²-value by the sum of the partial R²-values for both alleles of the contributing parent, yielding “normalized partial R²-values”, hereafter referred to as empirical genotype probabilities. If a contributing parent only featured a single allele, the probability of this allele automatically equals one.

#### 5.3.4 Breakpoint detection for haploblock definition

Gene level based genotypes still include artefacts or missing data, also due to filtering or other preprocessing steps mentioned higher. Hence, additional genotype clean-up is essential and was performed by the delineation of haploblocks. Importantly, for each part of the genome, up to two haploblocks can be identified per founder, in line with their (potentially) heterozygous nature. Essentially, for each parent, we infer haploblocks through linkage by evaluating those regions in mapping population individuals that are composed from a haplotype of the parent of interest and a haplotype from any other parent.

Specifically, within a sliding window of 30 genes (with a step size of 10 genes), we selected all mapping population samples that originated – within that region – from the chosen parent and any other parent, and obtained empirical genotype probabilities (see Methods section 5.3.3) for the alleles of the selected parent. When considering two selected samples, the alleles inherited from the same parent under study will either be consistently the same or consistently different, unless recombination has occurred. Therefore, to identify recombination events, i.e. haploblock edges, each pair of selected samples was considered and for the DNA segments from the shared parent the matching probability for each gene calculated, i.e. the chance that two selected samples inherited the same allele from the chosen parent. The matching probability was calculated using the following formula:

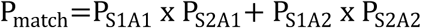

Where P_match_ is the matching probability of two samples, S_1_ and S_2_; P_S1A1_ is the empirical genotype probability for parental allele A_1_ in sample S_1_; P_S2A1_ is the same for sample S_2_; and similarly, P_S1A2_ and P_S2A2_ are the empirical genotype probabilities for parental allele A_2_ in samples S_1_ and S_2_, respectively. If samples had missing values for the empirical genotype probability for certain genes, the matching probability was set to 0.5. For each pair of samples, this results in a stretch of matching probabilities over the genes from the same parent under study.

These stretches of matching probabilities were subsequently used as input for the Segmentr R package (v0.2.0)^39^ to identify breakpoints in the founder haplotypes (“exact” option). These breakpoints correspond to recombination events/haploblock edges, where shared founder alleles shift to different founder alleles or vice versa. The package identified breakpoints in each stretch by maximizing the following function, modified from the documentation of the package:

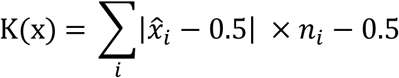

Where *x_i_* = {*x*_1_, *x*_2_, … , *x_n_* } are the matching probabilities of identified substretch i with length n. Thus, this function is maximized when the sequence is split into stretches that are homogeneously 0 or 1. A final 0.5 is subtracted as it will penalize overly short stretches in a sense that an extra breakpoint will automatically lead to (at most) a gain of 0.5 in the score, which is here subtracted again. By assessing the detected breakpoints for all pairs of selected samples in a window, we can infer recombination within genomic regions that were inherited from a single founder. To further ensure robustness, we filtered identified breakpoints, retaining only those detected at least *s−1* times, where s is the number of samples in which the matching probability for the gene (i.e. the breakpoint) was considered. This choice of *s* is based on the idea that if a breakpoint is unique to a specific sample, it should appear in every pair that involves that sample. This procedure implies that genotyping errors for a given gene and sample will typically not lead to breakpoints, and hence handles both missing data and genotyping errors.

Based on the detected breakpoints, we defined the boundaries of homozygous and heterozygous genomic regions in the original founders. Therefore, we considered each parental line individually, dividing its genome in genomic regions defined by neighbouring breakpoints for that parent. Subsequently, per genomic region, samples where the parent under study contributed to, were identified, and mean and mean variance of the empirical genotype probabilities of both candidate alleles (or 1, when only a single parental allele was identified) for this founder estimated over all samples. If the mean empirical genotype probability was ≥ 0.90 and the mean variance was < 0.03 in such a parental region, we classified it as homozygous. Otherwise, it was considered heterozygous in the parental line, making further genotyping required for samples containing those parental alleles (see Methods section 5.3.5).

#### 5.3.5 PCA based genotyping

Though previous steps led to the identification of haploblocks, genotyping is still required for those haploblocks where at least one of the founding contributors was heterozygous. For a given parent, we randomly selected one of the two possible parental alleles, and collected all corresponding empirical genotype probabilities for all genes within the haploblock from all samples originating from that parent and any other parent. We then performed PCA on this *n x m* matrix (where *n* is the number of selected samples and *m* is the number of genes), which typically resulted in a clear separation of the two genotype groups along PC1. The PCA analysis was performed using the missMDA R package (v1.18)^40^, which handles missing values.

If PCA successfully separated the two genotype groups along the x-axis, we assigned samples with a PC1 value < 0 to one genotype group and those with a PC1 value > 0 to the other, reflecting consistently small or large allele probabilities for the selected allele. The major genotype group (i.e. containing most samples) was designated as group 1, the minor genotype group as group 2. For these groups, we calculated the mean empirical genotype probabilities for the selected alleles, serving as a quality check for the genotyping process. Successful genotyping was indicated by opposite means for each gene in the two groups: if genotype group 1 had a mean < 0.5 for the selected allele, genotype group 2 should have a mean > 0.5, and v.v.. If this criterion was not met for at least 65% of the genes in the bin, we did not use the PCA-based genotypes for eQTL analysis. Instead, both alleles of that parent were collapsed into a single allele and further used throughout. Additionally, the mean empirical genotype probabilities for the selected alleles indicate which alleles per gene corresponded to each genotype group, which is crucial for accurately assigning genotype groups to samples originating from a single founder based on their corresponding alleles.

With the previous step, we could assign samples originating from the selected parent and any other parent to a specific genotype group, thus assigning a certain haploblock allele (a1 or a2). However, samples that inherited both alleles from the (same) selected parental line (2x a1, 2x a2, or a1 and a2) were thus far not included. To determine the haploblock alleles these samples inherited, we fitted a weighted linear regression model for each gene in the haploblock, similar to the one described in section 5.3.3. For each gene, the dependent variable was the allele fraction of the variant SNP allele, with the weights being the total counts at the SNP differing between the alleles of the two genotype groups. The independent variables indicated the presence (1) or absence (0) of the variant allele in the two previously identified genotype groups. The model was fitted three times: once with only the allele of genotype group 1, once with only the allele of genotype group 2, and once with the alleles of both genotype groups, representing the three possibilities for inherited alleles. The model with the highest total adjusted R² was selected. Each gene was then assigned a 0 (if the first model had the highest R²), 1 (if the second model had the highest R²), or 2 (if the third model had the highest R²). To assign the corresponding haploblock alleles, the values were averaged for all genes within the haploblock for that particular sample. If the average was closest to 0, the genotype was assigned as 2x a1 (homozygous); if the average was closest to 1, the genotype was assigned as a1 and a2 (heterozygous); if the average was closest to 2, the genotype was assigned as 2x a2 (homozygous).

#### 5.3.6 Genotype bins and variables

Finally, the union of the parental haploblock boundaries and inter-parental recombination events identified by the HMM was used to define bins, i.e. genomic regions in which each sample has a consistent genotype. These bins were subsequently translated into genotype variables for each parental line and possible allele, encoded as 0, 1 and 2, reflecting the number of copies present in the sample (Supplementary Figure 1).

Since the regularized regression approach (LASSO, see Methods section 5.4) cannot handle missing values in the input genotypes, any missing genotype variables were imputed using the mean dosage of the nearest neighboring non-missing genotypes. In addition, if a single homozygous bin appeared between heterozygous bins with identical sample genotypes, it was removed, and the heterozygous bins were merged.

### 5.4 eQTL mapping

#### 5.4.1 Single variant analysis

For the single-variant analysis, we employed MatrixEQTL (v2.3)^33^, considering associations significant if the FDR was < 0.01. To match the LASSO analysis, TMM normalized expression data was log_2_-transformed prior to fitting the linear models. To account for linkage, which can lead to adjacent significant bins near the actual most significant bin, we pruned the results based on linkage information. Specifically, for genotype variables originating from the same parental line and the same chromosome that are associated with the same target gene, we calculated the Pearson correlation between the genotypes. If a pair had a correlation ≥ 0.50, they were considered to be linked, and only the variable with the lowest p-value was retained.

#### 5.4.2 LASSO

We performed regularized regression analysis using a LASSO model implemented in the glmnet R package (v4.1.8)^41^. TMM-normalized expression data was log_2_-transformed and used as the dependent variable, while predictor variables included all parental genotype variables with a minor allele fraction of at least 0.05. The dataset was split into 4 folds, with each fold serving as a test set once, while the remaining 3 folds were used for training. To standardize the dependent variable, we subtracted the mean and divided by the standard deviation of the training data. The same scaling (based on the training set’s mean and standard deviation) was applied to the test set to avoid bias. For each fold, the LASSO model was trained on the standardized training data, with the optimal lambda parameter determined through additional 5-fold cross-validation. The model’s predictions, representing genetically determined gene expression, were obtained for the test data. We also recorded the bins selected as predictive features by the model. This process of 4-fold cross-validation was repeated 25 times to ensure robust results, yielding the mean genetically determined gene expression for each gene and sample across the 25 repetitions (based solely on the prediction of the test sets). For each gene, we calculated the R² value, which quantifies the proportion of variance in gene expression explained by the genotypes. This was done by fitting a linear model with the expression data as the dependent variable and the genetically determined gene expression as the independent variable. Genes were retained for eQTL selection if the R² was at least 0.01.

For eQTL selection, we first pruned the results within each fold to account for linkage. If multiple genotype variables from the same parental line and chromosome were selected in a given fold, we calculated their Pearson correlation. When the correlation was ≥ 0.70, only the variable with the highest absolute beta (LASSO-estimated coefficient) was retained. After linkage pruning within each fold, we merged the results across all 100 folds and counted the number of times each genotype variable was selected. We then performed a final round of linkage pruning on the combined results, retaining only the variable with highest selection count while merging the counts of linked variables (correlation ≥ 0.70). Results were only pruned or merged if the effects had consistent effect directions. Finally, genotype variables were classified as eQTLs for a gene if their (merged) selection count was at least 65.

This entire procedure was repeated independently for each minimally expressed gene using permuted data, for which the association is broken between genotypes and expression. The results from the permuted data were used to estimate the number of false positives corresponding to a certain selection count threshold.

#### 5.4.3 *Cis*- and *trans*-eQTL classification

To distinguish between *cis*- and *trans*-eQTLs, we analyzed the distribution of distances between eQTL and target gene (within 2 Mb from target genes), which are expected to be highly enriched for *cis*-eQTLs. We found that the background level was reached at approximately 1 Mb (Supplementary Figure 5). Consequently, we categorized associations within 1 Mb of their respective genes as *cis*-eQTLs, while those beyond this range were classified as *trans*-eQTLs. To identify *trans*-eQTL hotspots, we counted the number of *trans*-eQTLs in non-overlapping 300 kb windows across the genome. We initially selected windows that regulated at least 50 genes in *trans*, and merged adjacent selected windows. In the final selection, hotspot regions were defined as those affecting ≥ 75 genes in *trans*.

### 5.5 Association with phenotype

For each individual in the mapping population, we counted the number of eggs: (1) after 24 hours on their initial host, bean, (2) after 24 hours on the new host, tomato, and (3) after 48 hours on the new host. For one individual that laid no eggs on bean other fitness effects were expected and this sample was excluded from further analysis. The number of eggs laid between 24h and 48h on tomato was calculated by subtracting (2) from (3). For each of the 3 phenotypes, we fitted a linear regression model to predict the phenotype using the genetically determined expression (i.e. the average LASSO prediction) and extracted the corresponding p-value.

We subsequently categorized genes based on whether their expression was predominantly regulated in *cis* or *trans*. Therefore, we calculated the proportion of expression variance explained by *cis* and *trans* effects. First, eQTLs classified as *cis* or *trans* (see Methods section 5.4.3) were used as input of a multiple regression model (Anova, type II), and the partial eta² values (obtained from Effectsize R package, v0.8.9) summed over *cis-* resp. *trans*-eQTLs were used to estimate which fraction of the total R² from the LASSO model could be explained by *cis* resp. *trans* effects. Genes were subsequently classified into three groups: predominantly *cis* regulated (*cis* R² > 0.1, with *cis* effects explaining more than three times the variance of *trans* effects), predominantly *trans* regulated (*trans* R² > 0.1, with *trans* effects explaining more than three times the variance of *cis* effects), and mixed regulated genes (neither *cis* nor *trans* effects dominate but the total R² > 0.1).

For each of these groups we estimated the partial eta² for the effect of genetically determined expression on fecundity on tomato, adjusted for fecundity on bean, using the same procedure as above. Hence, we obtained an estimate for the variance in fecundity explained by the genetically determined expression. To enable the visual evaluation of statistical significance, we performed 1000 permutations per gene within each subgroup to generate null distributions. The resulting histograms of observed data hence also include density plots of the median trend under the null hypothesis, with 95% confidence intervals. The variance in fecundity explained by the genetic instrument, i.e. the partial eta², was also assessed for a group of genes where expression prediction was poor (total R² < 0.01), and no association with phenotype is expected. Within each subgroup, FDR values were obtained through Benjamini-Hochberg adjustment of p-values, and genes were deemed significantly associated with the phenotype if the FDR < 0.05. To identify *trans*-eQTL hotspots associated with phenotype, a Fisher exact test was used per hotspot to evaluate enrichment for *trans* target genes associated with phenotype among all predominantly *trans*-regulated target genes for that hotspot.

For the standard QTL approach, we fitted a linear model using the genotype as the independent variable and fecundity on tomato (between 24 and 48h) as the dependent variable while correcting for fecundity on bean, and extracted the corresponding p-value.

Narrow-sense heritability (h^2^) of uncorrected fecundity was estimated for each environment using a linear mixed model (LMM) implemented in the R package sommer (v4.3.4). Additive haplotype genotypes were used to construct a haplotype-based genomic relationship matrix (GRM; K = *ZZ*^T^/*m*). For each environment, we fitted the model *y* = *μ* + *a* + *e*, where the additive genetic effect was modeled as 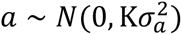 and the residual as 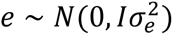. Variance components were obtained by REML, and narrow-sense heritability was calculated as 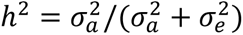.

### 5.6 Data processing and statistical analysis

All custom pipeline scripts and statistical analyses were implemented using Python (v3.10.6) and R (v4.2.1).

## Declarations

### Data availability

WGS data of the iso-female parents and the pooled sample of the population were previously deposited in the NCBI Sequence Read Archive (SRA) under BioProject PRJNA1170068. RNA-seq reads of the mapping population have been deposited in the NCBI SRA under BioProject PRJNA1225650. All code required to run the pipeline is available on Github (https://github.com/fedgraev/quasi_panmictic_eQTL).

### Authors’ contributions

TDM and TVL designed the experiment with input from RMC. BDB, FDG, SDR and MH performed the experiments. FDG and TDM designed and developed the bioinformatics pipeline with input from JF. FDG and TDM wrote the manuscript with contributions from TVL and RMC. All authors reviewed and approved the final manuscript.

## Acknowledgements

Not applicable.

