## Supplementary Figures for "Multiparental RNA-seq driven eQTL screening identifies loci underlying host plant fitness in a generalist herbivore"

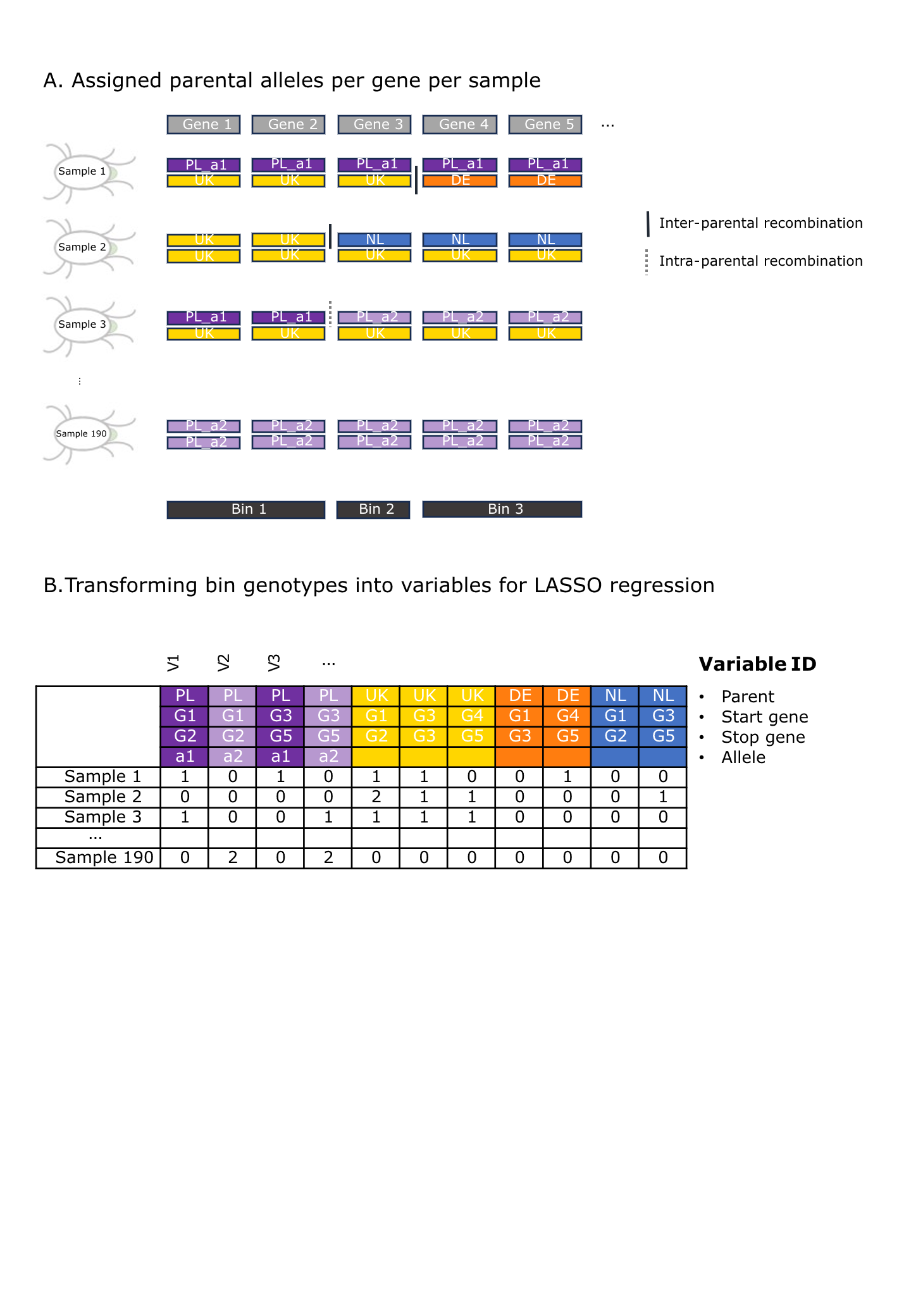
**Supplementary Figure 1. Overview of the binning approach**. This schematic (A) illustrates the binning approach using a simplified example with only 4 samples (out of 190 in the actual dataset) and 5 genes (out of ~18,000 in the actual dataset). The identified alleles per sample are shown, along with the inter- and intra-parental recombinations inferred from the HMM and haploblock delineation, respectively. This information is translated into genotype bins, which are subsequently transformed into variables for LASSO regression (B). These variables are parent-specific and defined by a start and stop gene. If a parent is heterozygous in a given region (e.g. PL in this example), the corresponding allele is also specified, hence a variable is generated for each allele.

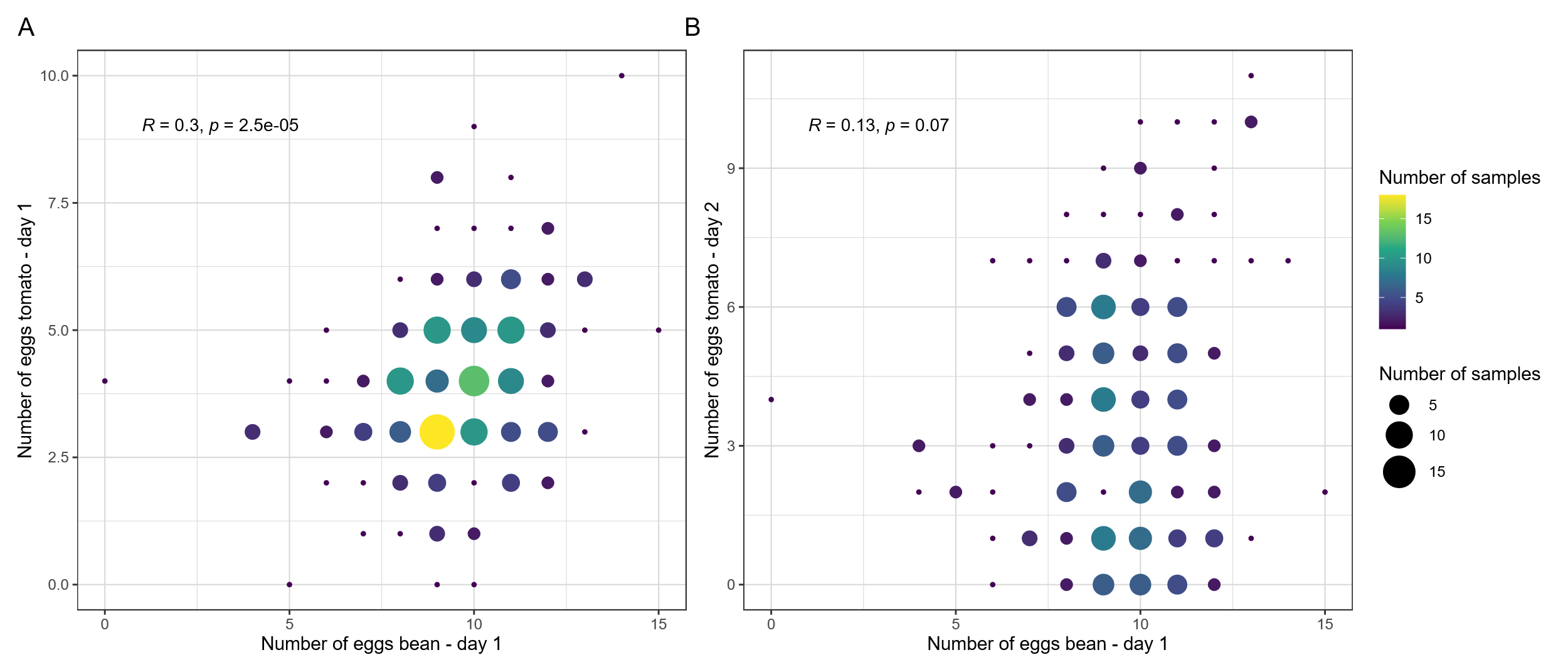

**Supplementary Figure 2. Relationship between the number of eggs laid by the mapping population individuals on bean and tomato.** Each point represents the exact number of eggs laid by a single female on bean on day 1 (x-axis) and on tomato (y-axis) on day 1 (A) or day 2 (B). Point size and color indicate the number of individuals with identical combinations of values. Gridlines correspond to discrete egg counts. Pearson’s correlation coefficient (R) and associated p-value are shown in each panel, quantifying the strength and significance of the relationship between fecundity on the two host plants.

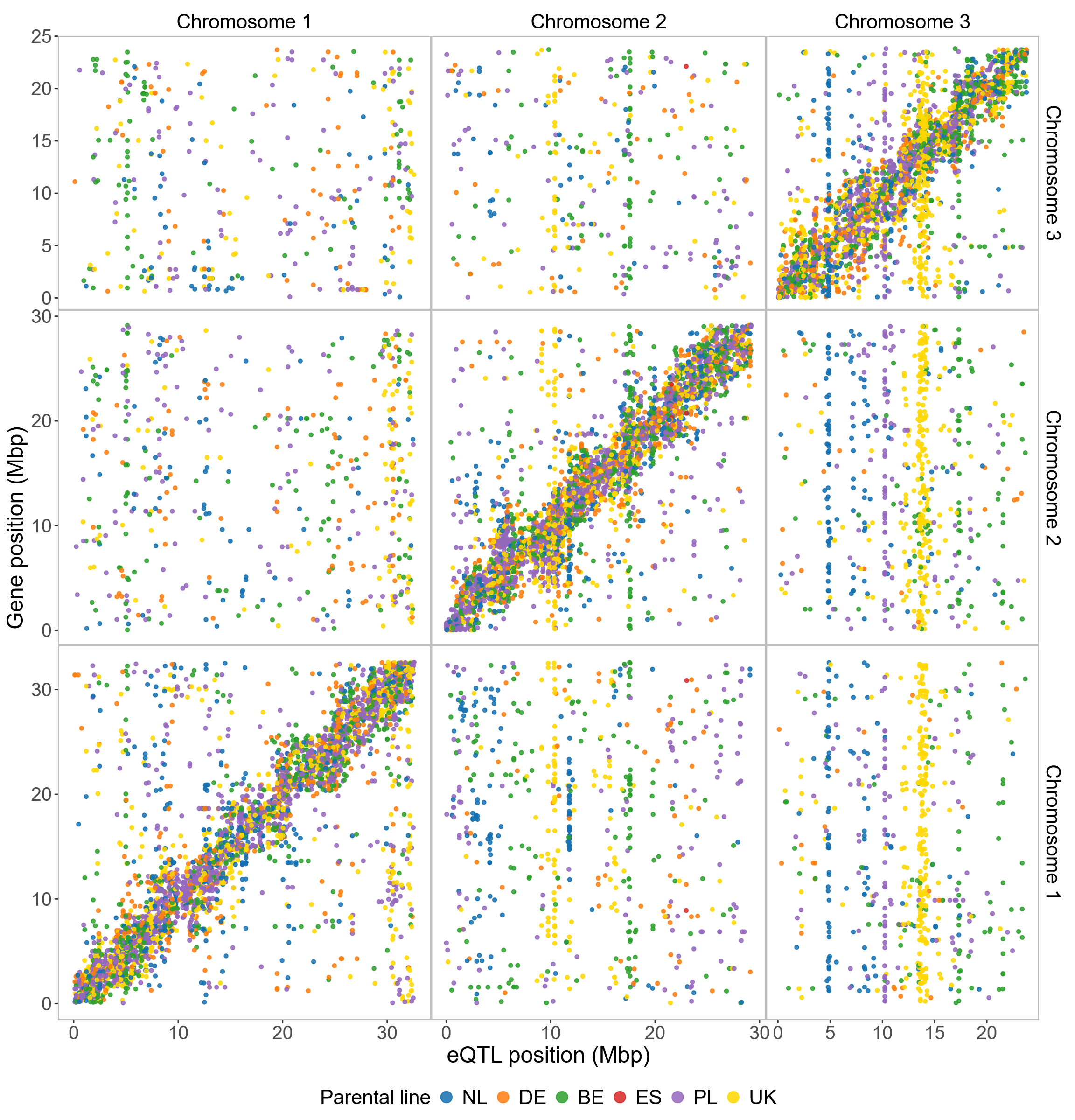

**Supplementary Figure 3**. **Significant eQTLs in a *quasi*-panmictic, multiparental mapping population based on single variant linear regression and comparison with LASSO results.** Scatter plot showing significant eQTLs (FDR < 0.01; 23,178 eQTLs in 8,273 genes on the 3 chromosomes) and the corresponding position of the eQTL (x-axis) and target gene (y-axis). Colors indicate the parental line of the eQTL as shown in the legend.

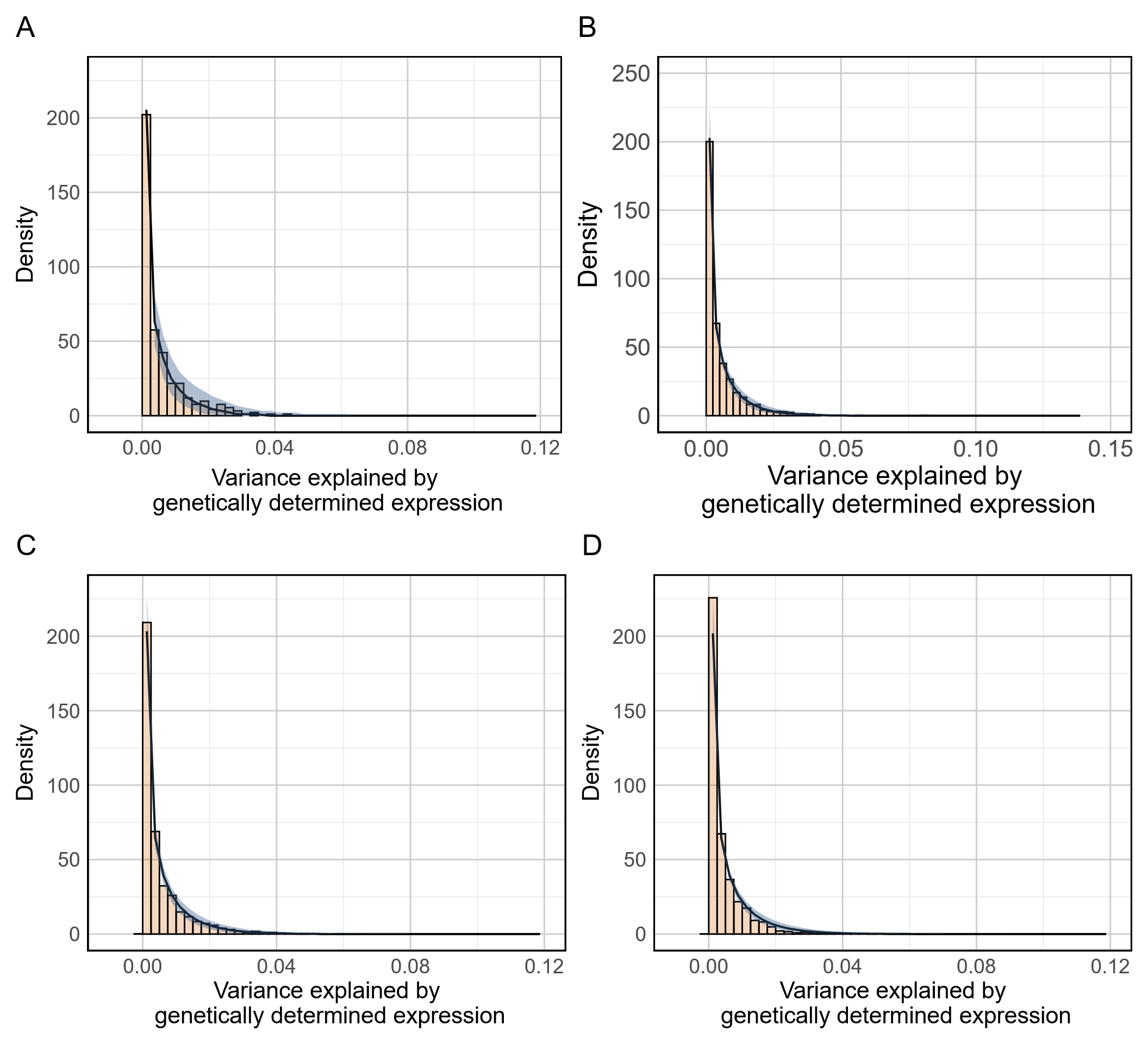

**Supplementary Figure 4**. **Histograms showing the variance in fecundity on the challenging host tomato during the first 24h explained by genetically determined expression.** The observed data are displayed as histograms, with the black density line representing the median trend under the null hypothesis and the blue shaded area indicating the 95% confidence interval. Subgroups are defined based on gene regulation: A) Predominantly *trans* regulated genes (n = 368), B) Genes with mixed regulation (n = 2,546) C) Predominantly *cis* regulated genes (n = 2,237), D) Genes with limited genetically determined gene expression (n = 4,063).

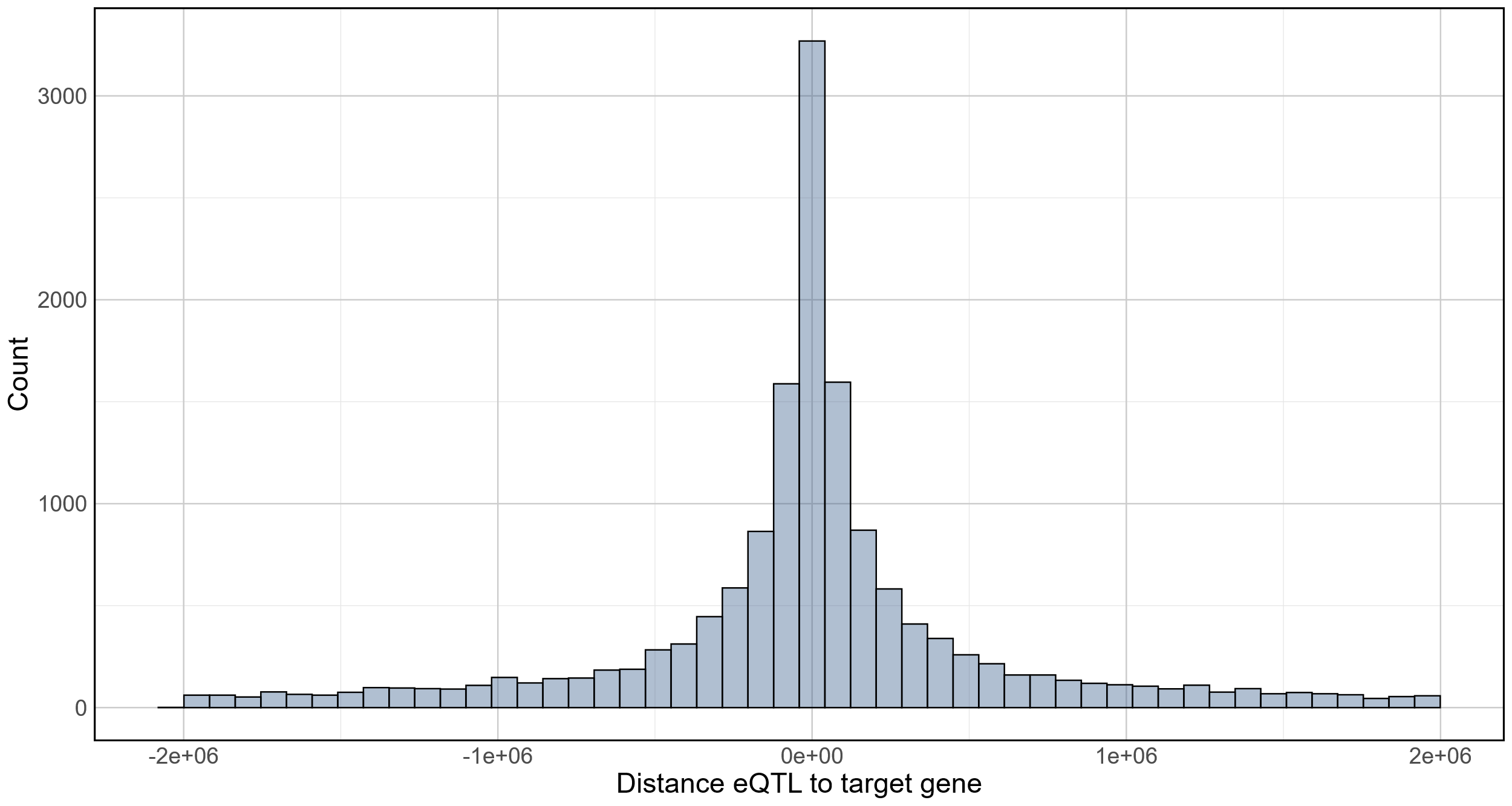
**Supplementary Figure 5**. **Histogram of the distances between eQTL and target gene**. Subset of genes where distances between eQTL and target gene are within 2 Mb.
