## Supplementary Methods for "Multiparental RNA-seq driven eQTL screening identifies loci underlying host plant fitness in a generalist herbivore"

### 1. Whole-genome sequencing of iso-female lines and the mapping population

Genomic DNA (gDNA) was retrieved from a sample of 400-800 mites per strain by a chloroform-phenol extraction as previously described^1^. Quality and quantity of the gDNA samples were assessed using a Denovix DS-11 spectrophotometer (DeNovix, Willmington, DE, USA) and by running a 2% agarose gel electrophoresis (30 min at 100 V). Sequencing libraries for the iso-female lines and a pooled sample of the mapping population (required to estimate the contribution of each strain to the final mapping population) were constructed using the NEBNext Ultra II Library Prep Kit (PCR-free) for Illumina at the NXTGNT sequencing facility (Ghent University, Ghent, Belgium), followed by sequencing at Genewiz (Leipzig, Germany) on an Illumina Hiseq3000 platform to generate 150 bp paired-end reads for the iso-female lines, and sequencing at NXTGNT (Ghent University, Ghent, Belgium) on an Illumina Nextseq500 platform to generate 75 bp paired-end reads for the mapping population.

All DNA reads were aligned to the London reference genome^2^ using BWA (v0.7.17-r1188)^3^ with default settings. The resulting BAM files were sorted by coordinates using SAMtools (v1.6)^4^, and duplicates were marked with Picard (v2.25.0) (https://broadinstintute.github.io/picard). Genetic variants were jointly called across all samples, following GATK’s (v4.2.0.0) best practices workflow^5^, and subsequently filtered according to GATK’s recommended criteria for hard filtering of germline variants: 1) MQ ≥ 40, 2) QD ≥ 2, 3) Q ≥ 100, 4) SOR ≤ 3, 5) FS ≤ 60, 6) MQRankSum ≥ -12.5, 7) ReadPosRankSum ≥ -8. Unique homozygous parental marker SNPs were identified. Based on the allele frequencies of these unique marker SNPs in the sequenced, pooled sample of the mapping population, we estimated the contribution of each parental line to this population, as previously described^6^.

### 2. RNA sequencing and phenotyping of single female mites

Total RNA was extracted from the frozen samples using the RNeasy Plus Mini Kit (Qiagen, Germany) according to the manufacturer’s Quick-Start Protocol and stored at -80 °C. The quality and quantity of the extracted RNA were analysed by NXTGNT (Ghent University, Ghent, Belgium) using the Bioanalyzer 2100 with the RNA 6000 Pico assay (Agilent Technologies, USA). For 190 samples with the highest RNA concentration and RNA integrity number (RIN) values, sequencing libraries were constructed using the Illumina Stranded mRNA Ligation Kit (Illumina, USA) at the NXTGNT sequencing facility (Ghent University, Ghent, Belgium), followed by sequencing at Genewiz (Leipzig, Germany) on an Illumina NovaSeq6000 platform to generate 150 bp paired-end reads.

The RNA-seq reads were aligned to the London reference genome^2^ using GSNAP (v2021-08-25)^7^, which performs variant tolerant mapping to reduce alignment bias. Known splice-sites were supplied using the GFF annotation described by Ji et al.^8^. Known variants were supplied as the VCF file generated from the WGS data of the iso-female founder strains (see Supplementary Methods 1). We selected the uniquely mapped reads and sorted the resulting BAM files by chromosomal coordinates prior to gene-level read counting with HTSeq (v2.0.2)^9^, accounting for strand-specificity and using the above-mentioned annotation. The count data was normalized using the edgeR (v3.42.4) TMM algorithm^10^, and genes were filtered based on minimal expression levels across all samples, dropping genes with raw read counts <10 in more than 171 samples (~90% of the samples). Additionally, we used the sorted BAM files to call genetic variants following GATK’s (v4.2.0.0)^5^ best practices workflow for RNA-seq to retrieve genotype probabilities. Although GATK recommends per-sample calling for RNA, this approach only identifies variable sites, preventing the distinction between homozygous reference genotypes and missing data. Since Brouard et al.^11^ validated the use of joint genotyping for RNA, we elected to call variants jointly on all samples.

As a proxy for the performance of female mites on a certain host, we counted the eggs laid per female every 24h. To facilitate this counting, leaf disks were photographed with a Nikon D5600 DSLR camera equipped with an AF-S Micro 60mm f/2.8 G ED macro lens. For a selection of images, we manually labelled the eggs and used sliced images from this set as a training set to train the YOLOv5 tool to automatically detect spider mite eggs^12,13^. To improve model performance, given the substantial differences in morphology of bean and tomato leaves, we trained separate models for each leaf type. Precision and recall, calculated on a validation image set, both exceeded 0.99, highlighting the excellent performance of the models.

### 3. Quality control

To evaluate the accuracy of our RNA-based genotyping strategy, we categorized bi-allelic SNPs into groups based on the level of mismatch between GATK- and haploblock-based genotyping results. SNPs were included in the analysis only if genotypes could be compared for at least 100 samples, meaning both approaches provided non-missing SNP genotypes for at least 100 samples. The SNPs were grouped based on the number of samples with mismatching genotypes: (i) no mismatches, (ii) more than 5 mismatches, (iii) more than 10 mismatches, and (iv) more than 25 mismatches. For all SNPs within each group, we calculated the inbreeding coefficient for both genotyping approaches.

### 4 Detoxification gene set

The set of detoxification genes, containing *T. urticae* gene families associated with xenobiotic metabolism, binding, or transport, was obtained from Ji et al^8^ and includes: ATP-binding cassette transporters (ABCs), cytochrome P450 monooxygenases (P450s), carboxyl-choline esterases (CCEs), major facilitator superfamily (MFS) transporters, intradiol-ring cleavage dioxygenases (DOGs), UDP-glucuronosyltransferases (UGTs), lipocalins, short-chain dehydrogenases (SDRs), Glutathione S-transferases (GSTs) and Lipase/lipooxygenases (PLATs).

### 5. Mapping population characteristics

Inter-parental recombination events were calculated based on shifts in parental origin as determined by the HMM. This allowed us to calculate the average number of inter-parental recombination events per sample and per chromosome. However, these inter-parental recombination events underestimate the total recombination events because they do not account for intra-parental recombinations. To estimate the total number of recombination events, assuming no inbreeding and a stable contribution of each parental line, we calculated the probability that a mapping population sample originates solely from one parental line as:

$$P_{hom}= \sum_{i} p_{i}^{2}$$

where p_i_ is the proportion contributed by the i-th parental line. The probability that a sample originates from two different parental lines is then 1 - P_hom_. Hence, we estimated the total number of recombination events as the observed number of inter-parental recombination events divided by 1 - P_hom_.

### 6. Gene ontology (GO) enrichment analysis

GO enrichment analyses were performed with the ‘enricher’ function of the package clusterProfiler in R (v4.8.3)^36^, using a BH-adjusted p-value cut-off of 0.05. The background gene set for GO enrichment analysis of *trans* hotspot targets consisted of the genes with a minimal expression level (see Supplementary Methods section 2) that were included for eQTL mapping. The Molecular Function (MF) and Biological Process (BP) terms were sourced from the ORCAE database (v01252019)^37^.
